# Metabolic regulation and structural mechanism of glutamine synthetase AMPylation

**DOI:** 10.64898/2026.09.02.748916

**Authors:** Eduardo Sabatine Lopes, Eijaz Ahmed Bhat, Larissa Fonseca Tomazini, Adriano Alves Stefanello, Lin Zhang, Paulo Sérgio Bueno, Oliver Einsle, Emanuel Maltempi de Souza, Edileusa Cristina Marques Gerhardt, Marco Aurelio Schuler de Oliveira, Khaled A. Selim

**Author notes:** Corresponding authors: Marco Aurelio Schuler de Oliveira. Avenida Colombo, 5790. Maringá-PR. Brazil. CEP 87020-900.; Khaled A. Selim. Microbial Biochemistry Group, Institute of Phototrophic Microbiology, Heinrich-Heine University Düsseldorf, Düsseldorf 40225, Germany. These authors contributed equally (Eduardo Sabatine Lopes, Eijaz Ahmed Bhat, Marco Aurelio Schuler de Oliveira, Khaled A. Selim).

## Abstract

In bacteria, glutamine synthetase (GS) is the main ammonium assimilation enzyme. Its activity is tightly regulated according to cellular energy status and carbon–nitrogen balance through reversible AMPylation catalyzed by the bifunctional enzyme GlnE, which is controlled by the signal transducer GlnB protein. Although GS AMPylation has been extensively studied, the GlnB:GlnE:GS regulatory pathway exhibits substantial plasticity among bacterial groups, and the structural basis of GS inhibition by AMPylation remains unclear. Here, we describe how carbon, nitrogen, and energy signals regulate GS AMPylation in *Herbaspirillum seropedicae* and uncover the structural mechanism underlying enzyme inhibition. Our data reveal that GS AMPylation is independent of unmodified GlnB, whereas uridylylated GlnB (GlnB-UMP) inhibits AMPylation under nitrogen-limiting conditions through a GlnB-GlnE complex modulated by 2-oxoglutarate. We further show that GlnE directly senses glutamine under nitrogen-sufficient conditions, with signal integration depending primarily on energy availability. To elucidate the mechanism of AMPylation GS inhibition, we solved Cryo-EM structures of unmodified and AMPylated GS in complex with MgATP and MnADP. Structural comparisons revealed that AMPylation increases the flexibility of the AMP-loop, disrupting a hydrogen-bond network that stabilizes Arg342 in the orientation required to position the ATP γ-phosphate in its catalytic conformation for efficient phosphoryl transfer to glutamate. These findings reveal how metabolic signals are integrated to regulate GS AMPylation and provide the first structural insights into the mechanism underlying bacterial GS inhibition by AMPylation.

**Importance:** Glutamine synthetase is the central enzyme of bacterial nitrogen assimilation and has been studied for more than six decades. Although its regulation by reversible AMPylation has long been recognized, the structure of the AMPylated enzyme has remained unknown, leaving the molecular basis of inhibition unresolved. We present the first structures of AMPylated glutamine synthetase, revealing that the modification increases the flexibility of a surface loop that disrupts the positioning of a catalytic residue required for ATP utilization. Together with the identification of an alternative signaling mechanism controlling AMPylation in *Herbaspirillum seropedicae*, these findings resolve a longstanding question in bacterial nitrogen metabolism and uncover unexpected diversity in its regulation.

## Introduction

In bacteria, glutamine and glutamate are the main nitrogen donors for bacterial anabolism, since they are involved in several transamination reactions, such as in the biosynthesis of amino acids, nucleotides, and others. Under nitrogen-limiting conditions, glutamine and glutamate are produced by glutamine synthetase (GS, EC 6.3.1.2) and glutamate synthase (GOGAT, EC 1.4.1.14) enzymes in the GS-GOGAT cycle that acts as a metabolic hub, controlling the homeostasis of glutamine and glutamate—which are vital for cellular viability (Reitzer, L. 2003; Forchhammer & Selim 2020).

The regulation of the GS-GOGAT cycle is mainly through the control of GS activity. In nature, the GS regulation is variable. Bacterial GS is regulated at the transcriptional level and at the protein level through post-translational modification and allosteric regulation. Other mechanisms, such as assembly modulation (Schumacher, M.A. et al., 2023) and protein-protein interaction (Herdering, E. et al. 2025), have been described for Archaea. In *Escherichia coli,* GS is allosterically inhibited by the end products of glutamine metabolism, glycine, and alanine. This negative feedback is cumulative, meaning that each modulator has only a partial effect on GS activity, but their effects are additive when they occur simultaneously. In *Herbaspirillum seropedicae*, the target organism in this work, GS is mainly inhibited by threonine and tryptophan (Tomazini, L.F. et al., 2025). Transcriptionally, GS is controlled via the Ntr System, the master regulator of nitrogen metabolism that responses to nitrogen-carbon balance and energy availability. PII proteins, encoded by *glnB*, are central components of the Ntr System. PII proteins are homotrimers that regulate several cellular processes by protein-protein interaction (Selim & Alva 2024; Forchhammer et al. 2022). For instance, PII regulates key enzymes of nitrogen metabolism, such as the rate-limiting enzyme of arginine biosynthesis, N-acetyl-L-glutamate kinase (Selim et al., 2020a,b; Lapina et al., 2018), for which we recently elucidated the mechanism of PII regulation (Elshereef et al., 2026). The interaction between PII and its targets usually occurs through the PII T-loop, which assumes different conformations in response to the binding of ATP, ADP, and 2-oxoglutrate (2OG), a TCA cycle metabolite (Selim et al., 2019; Forchhammer & Selim 2020). Furthermore, the T-loop is subject to a reversible post-translational modification, uridylylation, carried out by the bifunctional enzyme GlnD (EC-2.7.7.59) in response to glutamine/2OG ratio (Emori, M.T. et al., 2018; Bonatto, A.C. et al., 2007). The PII modification and the binding of allosteric effectors drive the formation of PII-target complexes (Forchhammer et al. 2022; Huergo, L.F. et al, 2013).

Moreover, GS activity is modulated by a reversible post-translational modification — AMPylation — carried out by the bifunctional adenylyl-transferase/adenylyl-removing enzyme GlnE (EC-2.7.7.42). In *E. coli*, both GlnE activities are strictly dependent on PII. When nitrogen is available, unmodified PII forms a complex with GlnE, stimulating its adenylyl-transferase (AT) activity, which inhibits GS by the addition of an AMP moiety at a highly conserved tyrosine residue (Atkinson, M.R. & Ninfa, A.J., 1998; Jiang, P. & Ninfa, A.J., 2009; Jiang, P. et al., 2007). AT activity is also stimulated by the direct sensing of glutamine by GlnE (Jiang, P. et al., 2007). On the other hand, in nitrogen starvation, PII is uridylylated and forms a new complex with GlnE, promoting its adenylyl-removing (AR) activity, thereby leading to GS activation (Jiang, P. et al., 2007). However, the molecular basis of GS inhibition by AMPylation remains elusive.

Despite the well-understood GlnE regulation in *E. coli*, it seems that its regulatory mechanism is not conserved in the Bacteria domain. In *Rhodospirillum rubrum*, an Alphaproteobacterium, GlnE is neither regulated by uridylylated PII nor glutamine. Its AT activity is activated by interaction with unmodified PII, and the complex formation is modulated by the 2OG levels that prevent the complex formation (Jonsson, A. et al, 2007). The *R. rubrum* GlnE AR activity seems to be constitutive (Jonsson, A. et al, 2007). The GS AMPylation is not restricted to Proteobacteria, but it is also found in Actinobacteria, such as Gram-positive *Streptomyces coelicolor*. In *S. coelicolor*, however, the GlnE regulation is completely independent of PII and GlnD, since the GS regulation was not affected in both PII and GlnD mutants (Fink, D. et al., 1999). Another difference is that the PII homolog GlnK is adenylylated by GlnD instead of uridylylated, and then GlnD is an adenylyltransferase (Hesketh, A}. et al, 2002).

Understanding the GlnB-GlnE-GS regulatory cascade has biotechnological importance, as it enables the development of endophytic nitrogen-fixing bacteria engineered to fix nitrogen and release excess ammonium to the host plant —a strategy recently achieved in a similar PII-regulated system (Gerhardt & Selim 2026; Batista et al. 2025). Here, we described the regulation of the PII-GlnE-GS cascade by the carbon-nitrogen balance signals in the Betaproteobacteria *Herbaspirillum seropedicae* SmR1. *H. seropedicae* is an endophytic nitrogen-fixing bacterium found associated with economically important Poaceae plants, such as rice, sugarcane, maize, and sorghum (Baldani, J.I. et al., 1986; Muthukumarasamy, R. et al., 2006). Notably, *H. seropedicae* could be genetically engineered and employed as a biofertilizer to enhance the incorporation of fixed nitrogen into plant biomass (Pankievicz, V.C. et al., 2015).

*H. seropedicae* has two PII homologous proteins, encoded by the *glnB* and *glnK* genes (Noindorf, L. et al., 2006; Benelli, E.M. et al., 1997). Here, we demonstrate that uridylylated GlnB transduces carbon-nitrogen balance status by inhibiting GS AMPylation through the formation of an inhibitory GlnB-UMP:GlnE complex that is modulated by 2OG levels. On the other hand, the GS AMPylation is GlnB-independent and occurs in response to glutamine levels, sensed directly by GlnE. Using single particle cryo-EM, we solved the structures of unmodified GS and AMPylated GS, revealing the precise molecular and structural mechanisms of GS inhibition by AMPylation. We demonstrated that addition of an AMP-moiety to the conserved AMPylation residue Tyr400 increases the flexibility of GS AMP-loop, affecting nucleotide binding in the active site and thereby explaining the molecular details of AMPylation-dependent GS inhibition. Ultimately, our study advances our understanding of PII regulatory plasticity and uncover the molecular mechanism of GS AMPylation in *H. seropedicae*.

## Results

### *H. seropedicae* GS (HsGS) kinetics: *in vivo* switch-off and *in vitro* AMPylation

The GS AMPylation state changes in response to the ammonium availability, thereby controlling the enzyme activity according to the cell’s metabolic needs. To investigate the AMPylation control on HsGS activity, the enzyme was first overexpressed and purified to high homogeneity (figure S1). The kinetics properties of GS were determined *in vitro* through the GS PK/LDH dehydrogenase assay. As shown in figures 1A and 1B, the K_M_ for NH_4_^+^ was determined as 0.15 mM and, for glutamate as 0.80 mM.

**Figure 1.**
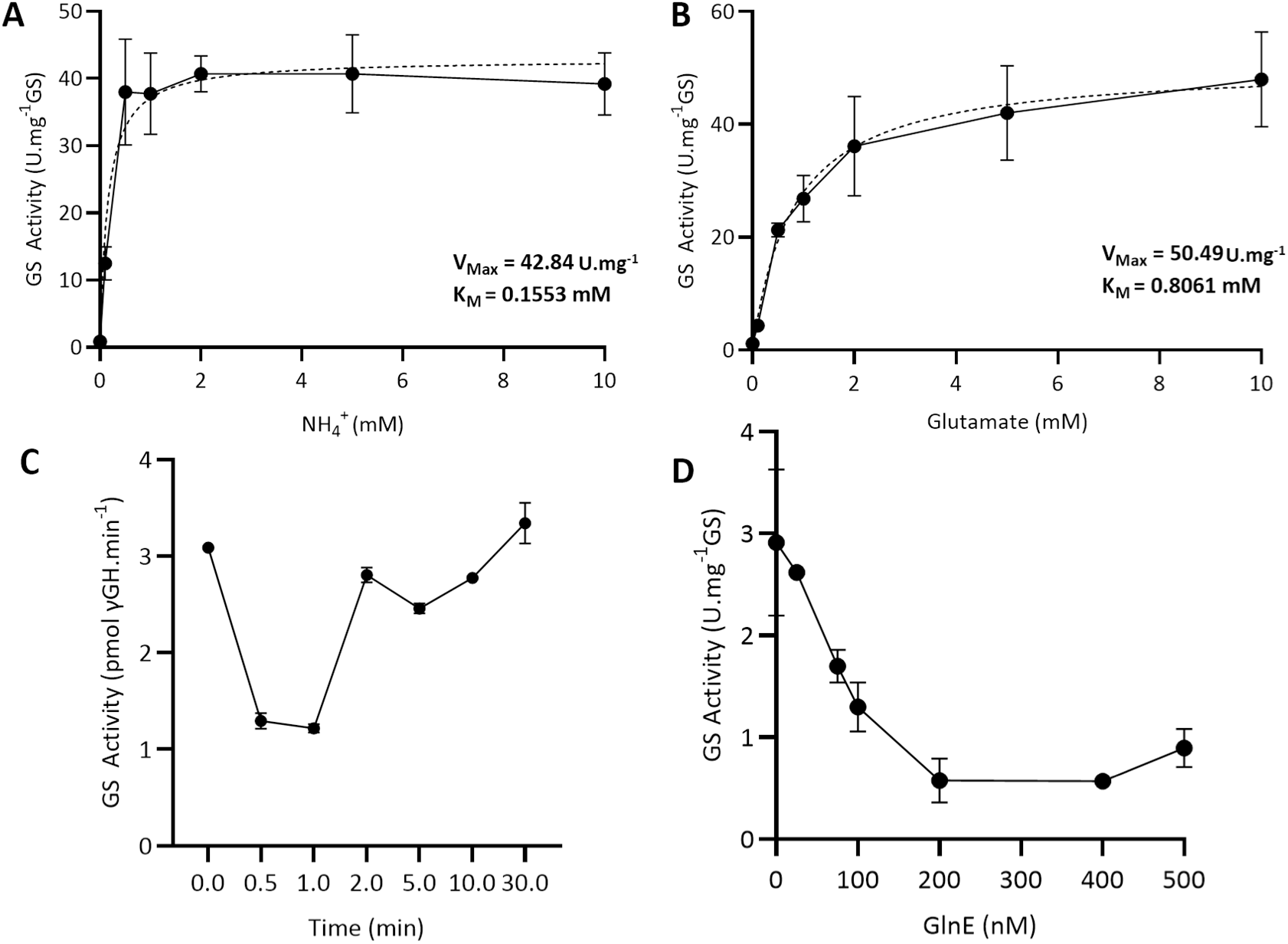
*in vitro* and *in vivo* HsGS activity. A and B) HsGS kinetics for NH_4_^+^ and glutamate using PK/LDH coupled assay. The K_M_ for NH_4_^+^ was determined as 0.15 mM and, for glutamate as 0.80 mM, while the V_Max_ was comparable for both. C) *In vivo* GS switch-off. The biosynthetic GS activity dropped directly after the addition of 200 µM NH_4_Cl until its consumption, indicating the GS switch-off through the transition from an unmodified to a modified state. For the GS switch-off, *H. seropedicae* cells grew in NFbHP-malate medium under nitrogen-starving conditions for 24 h before the addition of 200 μM NH_4_Cl. After the ammonium shock, aliquots were collected and immediately frozen in liquid nitrogen. The crude extract was then used to determine the *in vivo* GS activity. D) GlnE-dependent GS inhibition via the adenylyl-transferase activity. Purified GlnE inhibited GS activity in the PK/LDH assay, regardless of the presence of GlnB. The GS activity was carried out in triplicate, and the results are described as the mean ± SD.

To check whether the NH_4_^+^ availability could lead to a GS switch-off *in vivo*, we performed an ammonium-shock experiment by adding 200 μM of NH_4_Cl to nitrogen-starved cultures. The HsGS was inhibited by 3-fold already after 30 secs of the NH_4_^+^ shock, and maintained that inhibition for a min, suggesting a very fast response to modulate GS activity (figure 1 C). Two minutes after the ammonium shock, the GS activity was regained, suggesting ammonium consumption. This result indicates the GS switch-off through the transition from an unmodified state to the adenylylated state.

To follow *in vivo* GS inhibition via GlnE-dependent AMPylation, we performed *in vitro* GS inhibition assay in the presence of GlnE enzyme. For this purpose, we purified the HsGlnE to high homogeneity (figure S2). The GlnE molecular mass was determined as 93 ± 15.4 kDa by mass photometry and size exclusion chromatography, corresponding to a monomer (theoretical mass 104 kDa) (figure S2). GS activity assay was inhibited in a GlnE-dependent manner, indicating successful GS AMPylation to tune its activity (figure 1D). Notably, these reactions were performed in the absence of GlnB protein, demonstrating that HsGlnE-mediated AMPylation is GlnB-independent, in contrast to *E. coli* system, which requires GlnB for GS AMPylation via GlnE.

### Glutamine negative feedback through GS AMPylation

In *E. coli*, the GS is not directly feedback-inhibited by its product, glutamine (Ree, et al., 1989); instead, the C-terminal adenylyl-transferase (AT) domain of GlnE acts as a glutamine sensor, enhancing GS AMPylation to suppress its activity in a glutamine-dependent manner (Jiang, P. et al., 2007). Conversely, GlnE-dependent GS AMPylation does not respond to glutamine in *R. rubrum*, suggesting that glutamine-mediated regulation via GlnE may not be a conserved feature (Jonsson, A. et al, 2007). To determine whether the HsGlnE C-terminal domain functions as a glutamine sensor, we generated an N-terminal truncated GlnE variant (ΔNT-GlnE; figure S3) comprising amino acids 438-928, which encompasses both the central domain and the C-terminal AT domain. As observed in *E. coli*, HsGS activity was not directly feedback-inhibited by glutamine (figure 2A,B). With ΔNT-GlnE alone, HsGS activity was reduced by half without exogenous glutamine addition and was completely abolished upon its addition in a dose-dependent manner (figure 2B). In absence of external glutamine, GS inhibition via GlnE-dependent AMPylation is likely driven by the glutamine produced via basal GS activity, an effect that intensified as external glutamine was added (figure 2B). Titration of glutamine into the GlnE-dependent AMPylation assay of GS revealed an estimated IC_50_ of 86.8 μM for glutamine, supporting high-affinity binding of glutamine to GlnE. Together, these results indicate the existence of a glutamine-sensory site on the HsGlnE C-terminal, as found in *E. coli*. As we did not observe a direct glutamine-dependent inhibition of GS, these findings suggest that the AMPylation serves instead as a mechanism for glutamine-mediated negative feedback in *H. seropedicae*. Consequently, sGlnE acts as a nitrogen sensor regulating the ammonium assimilation rate in *H. seropedicae*. In this context, the glutamine-dependent AMPylation lileky represents a conserved regulatory mechanism shared with *E. coli*, whereas its independence from GlnB does not.

**Figure 2.**
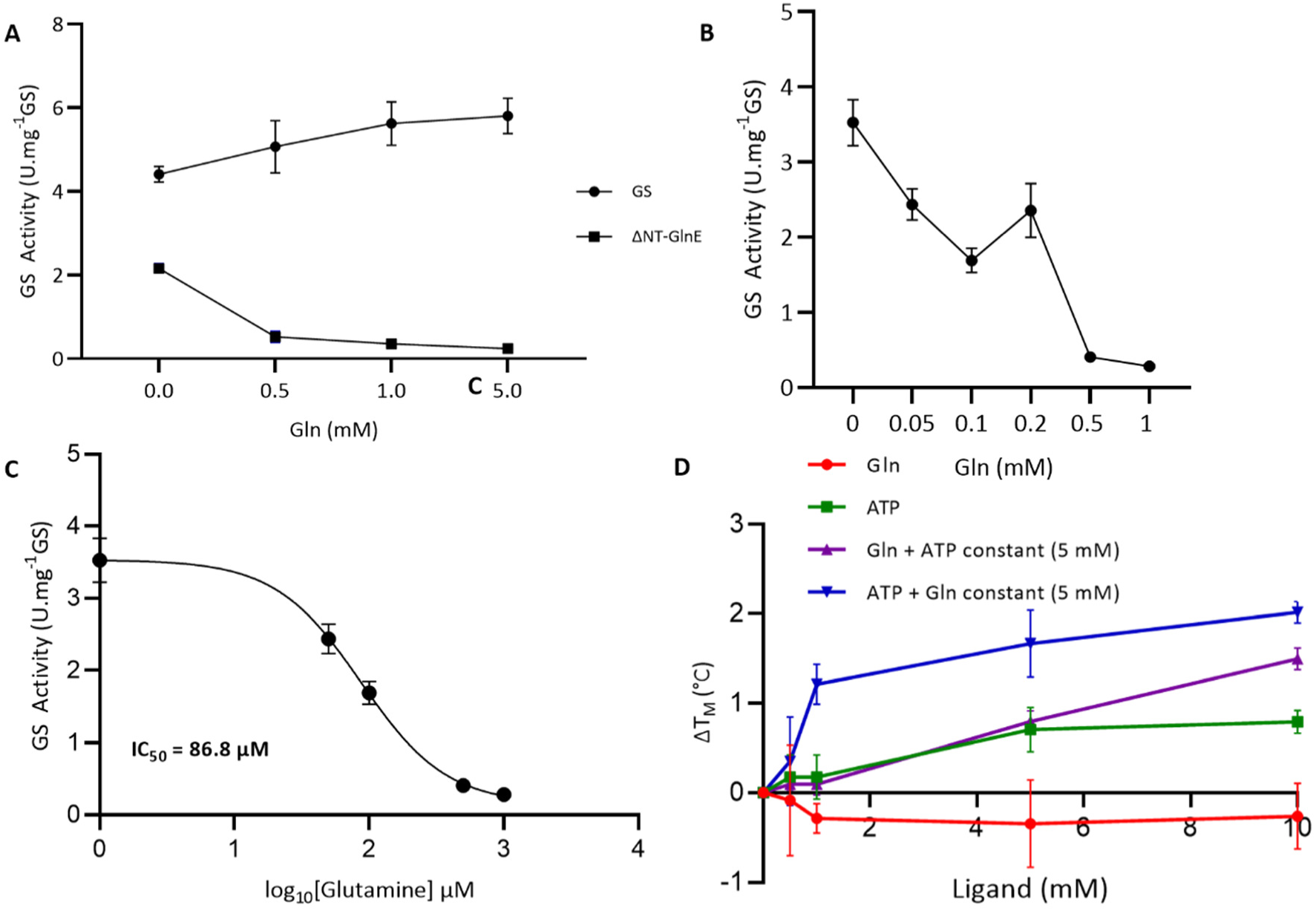
Glutamine negative feedback. The influence of glutamine on GS activity/AMPylation was monitored through the PK/LDH coupled assay. GS reaction was started by adding 0.5 mM glutamate (Glu) after 15 min of AMPylation reaction. The GS activity was not directly affected by glutamine, but it was reduced in the presence of ΔNT-GlnE and abolished when the reaction was supplemented with glutamine (A). The activity was inhibited by around 50% with the addition of 0.1 mM glutamine and abolished over 0.5 mM glutamine (B). C) The IC_50_ for glutamine was determined as 86.8 μM. D) Differential Scanning Fluorimetry (DSF) Analysis of Protein Stability in the presence of Glutamine and ATP. The graph shows the change in melting temperature (ΔT_M_) of a protein as a function of ligand concentration. The red line represents the effect of glutamine alone, showing no stabilization. The green line indicates the stabilizing effect of ATP alone, which increases with concentration and reaches saturation at 5 mM. The purple line demonstrates that glutamine induces further stabilization when ATP is present at a constant 5 mM. The blue line highlights a pronounced synergistic effect, where the presence of 5 mM glutamine significantly enhances the stabilizing effect of ATP, leading to the highest observed ΔTM. These results indicate a strong cooperative interaction between ATP and glutamine in stabilizing the protein. The experiments were carried out in triplicate, and the results are described as the mean ± SD.

To further characterize the glutamine-sensory domain of GlnE, we employed Differential Scanning Fluorimetry (DSF) to assess the thermal stabilization of the ΔNT-GlnE variant by ATP or glutamine. Titration of glutamine alone into ΔNT-GlnE did not induce thermal stabilization, as evidenced by the unchanged ΔT_M_ values (figure 2E, red line). In contrast, ATP titration increased the ΔT_M_ that reached saturation at 5 mM, indicating a direct stabilizing effect of ATP on ΔNT-GlnE (figure 2E, green line). To test the potential synergy between both molecules on ΔNT-GlnE stabilization, we performed titrations in the presence of a constant 5 mM concentration of the opposing ligand. In the presence of ATP, glutamine titration led to a concentration-dependent stabilization of the protein (figure 2E, purple line). Notably, titrating ATP in the presence of 5 mM glutamine also stabilized the protein, but saturation was achieved at a lower concentration of 1 mM (figure 2E, blue line). Together, our DSF data demonstrate a cooperative binding of ATP and glutamine to ΔNT-GlnE, hinting that ATP binds first, facilitating subsequent glutamine binding, with glutamine significantly enhancing ATP binding through positive cooperativity.

### GlnB and GlnB-UMP rules in GS AMPylation

GlnB proteins transduce nitrogen signals through protein-protein interactions, which are modulated by their post-translational modification status and by binding the allosteric effectors ATP, ADP, or 2OG (Forchhammer and Selim, 2020; Forchhammer et al., 2022). In *E. coli* and *R. rubrum*, unmodified GlnB is necessary to activate GlnE adenylyl-transferase activity. Conversely, our initial observations revealed that the HsGlnE activity is GlnB-independent (figure 1D). Nevertheless, to test whether GlnB or GlnB-UMP could modulate the GS AMPylation in the presence of allosteric effectors, we first purified the GlnB protein to high homogeneity and, where required, uridylyted it *in vitro* to prepare GlnB-UMP (figure S4).

In the presence of both GlnB and GlnE, the GS activity was strongly reduced, indicating successful adenylylation (figure 3A); however, a similar pattern was also observed in the absence of GlnB, supporting GlnB-independent GS adenylylation via GlnE. This AMPylation profile remained unchanged whether GlnB was saturated with ATP or ATP/2OG. GS activity was also inhibited by GlnE-dependent AMPylation in the presence of GlnB-UMP:ATP, but this inhibition was relieved when 2OG was added to the reaction (figure 3A), suggesting that the GlnB-UMP:ATP:2OG complex inhibits the GlnE adenylyl-transferase activity, thereby protecting GS from adenylylation.

**Figure 3.**
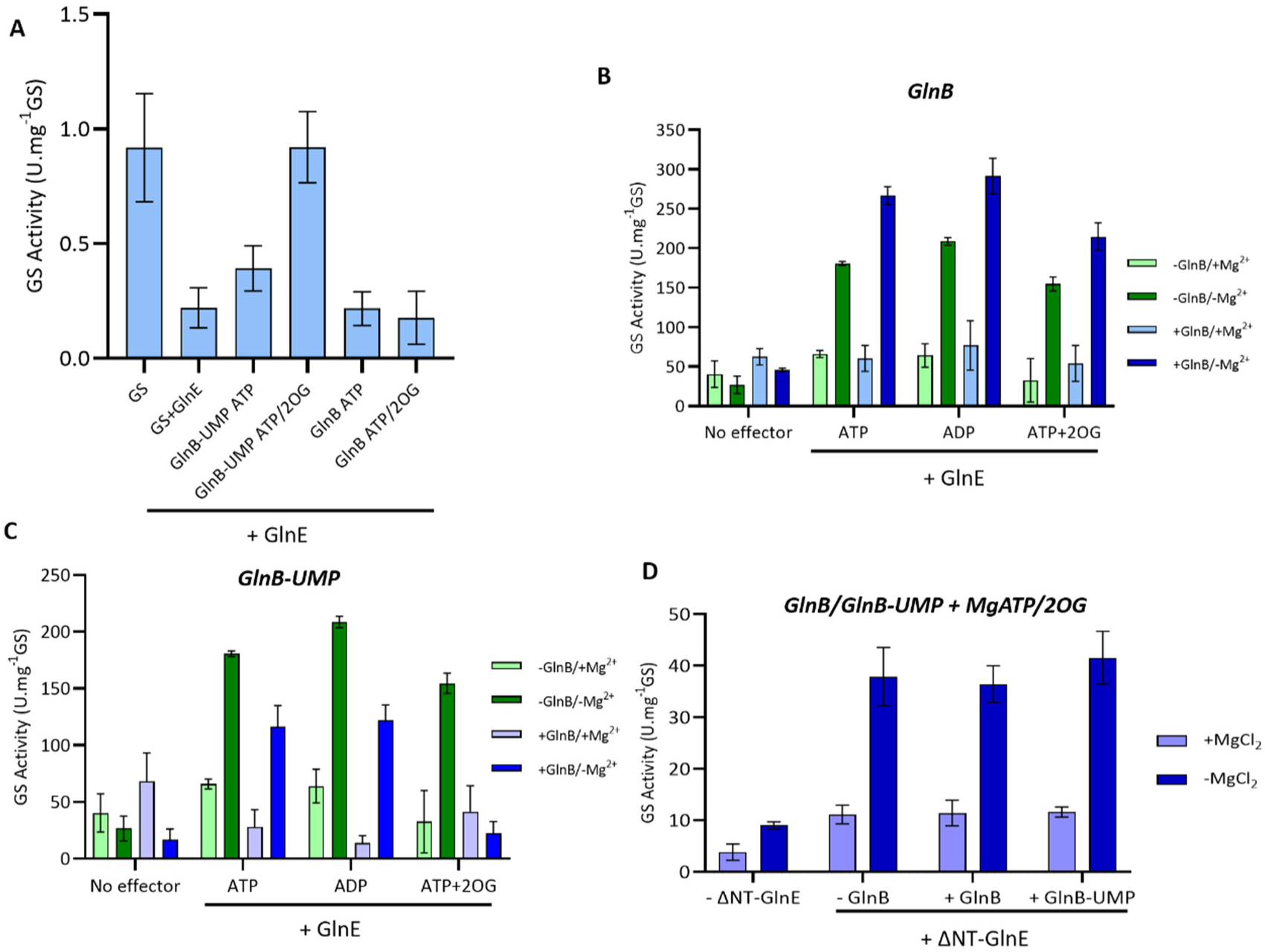
Regulation of GS AMPylation by GlnB and GlnB-UMP. A) PK/LDH-coupled GS activity assay showed inhibition of GS activity both in the presence and absence of GlnB+ATP, GlnB+ATP/2OG, and GlnB-UMP+ATP, indicating that neither GlnB nor GlnB-UMP had a regulatory effect on GS adenylylation. Conversely, the combination of GlnB-UMP+ATP/2OG inhibited GS adenylylation, resulting in activity levels similar to those observed without GlnE. With exception of GS alone control, all conditions were contained GlnE. B and C) γ-glutamyltransferase (γ-GT) assay corroborated the AMPylation pattern. Higher GS activity in the absence of MgCl_2_ compared to its presence indicated adenylylation. This AMPylation was inhibited by GlnB-UMP+ATP/2OG. No regulatory effect was observed in the presence of ADP. D) Regulation of the ΔNT-GlnE activity by GlnB and GlnB-UMP in presence of MgATP/2OG using the γ-GT assay. In all cases, the GS was adenylylated as evidenced by the higher GS activity in the absence of MgCl_2_, demonstrating that the inhibitory effect of GlnB-UMP was not observed on the truncated variant. The experiments were carried out in triplicate, and the results are described as the mean ± SD.

Since ADP interferes with the PK/LDH-based assay, the experiments were also carried out using the reverse γ-glutamyl-transferase (γ-GT) reaction of GS. In this assay, a higher GS activity in the absence of MgCl_2_ compared to that in the presence of MgCl_2_ indicates GlnE-dependent AMPylation (Tomazini, L.F., et al., 2025). In the γ-GT assay, GS was again adenylylated regardless of the presence of unmodified GlnB, even when ATP, ADP, or ATP/2OG were present in the reaction (figure 3B). Notably, GlnB-UMP:ATP and GlnB:ADP complexes did not affect GS adenylylation (figures 3B and 3C). However, GS AMPylation was once again inhibited when GlnB-UMP was present alongside ATP/2OG (figure 3C). These results indicate that AMPylation of *H.* seropedicae GS is regulated by a unique mechanism in which GlnB plays a distinct role, differing from the regulatory models previously described for *E. coli* and *R. rubrum*.

Following our characterization of wild-type GlnE regulation, we tested whether the truncated ΔNT-GlnE variant would respond similarly to GlnB-UMP-mediated inhibition. We used the γ-GT assay to assess GS activity and its AMPylation by ΔNT-GlnE in the presence of GlnB or GlnB-UMP combined with ATP/2OG. GS activity in the absence of MgCl_2_ remained higher than in its presence, indicating GS adenylylation under all conditions (figure 3D). This result validates our previous experiments, where unmodified GlnB:ATP:2OG failed to alter GlnE activity. On the other hand, the protective effect of GlnB-UMP:ATP:2OG complex, preventing AMPylation, was completely abolished. This lack of an inhibitory effect suggests that the N-terminal domain is essential for intramolecular signal transduction to the regulatory C-terminal domain of GlnE that could block the glutamine-sensitive site, the active site, or both. Notably, an analogues mechanism was previously shown for *E. coli*.

### 2OG modulation of GlnB-UMP:GlnE complex

As the inhibitory effect of GlnB-UMP on GlnE was observed in the presence of 2OG, we next titrated 2OG to determine the exact concentration required to form the GlnB-UMP:GlnE complex and inhibit AMPylation. Using the PK/LDH-based assay, we revealed that at least 1 mM 2OG is required to relieve GS inhibition, whereas GS remained inhibited at concentrations up to 0.5 mM of 2OG. This indicates that the transition between 0.5 and 1 mM 2OG triggers the GlnB-UMP:GlnE complex assembly (figure 4A). Next, we used mass photometry to characterize the 2OG-dependent GlnE:GlnB-UMP complex assembly. In the presence of 3 mM 2OG, we observed a molecular mass shift from 93 ± 15 kDa (GlnE alone) to 134 ± 23 kDa for the complex, indicating a 1:1 stoichiometry for the GlnE:GlnB-UMP complex (figures 4B and S5). Notably, no other tested conditions induced a mass shift suggestive of complex assembly (figure S5).

**Figure 4.**
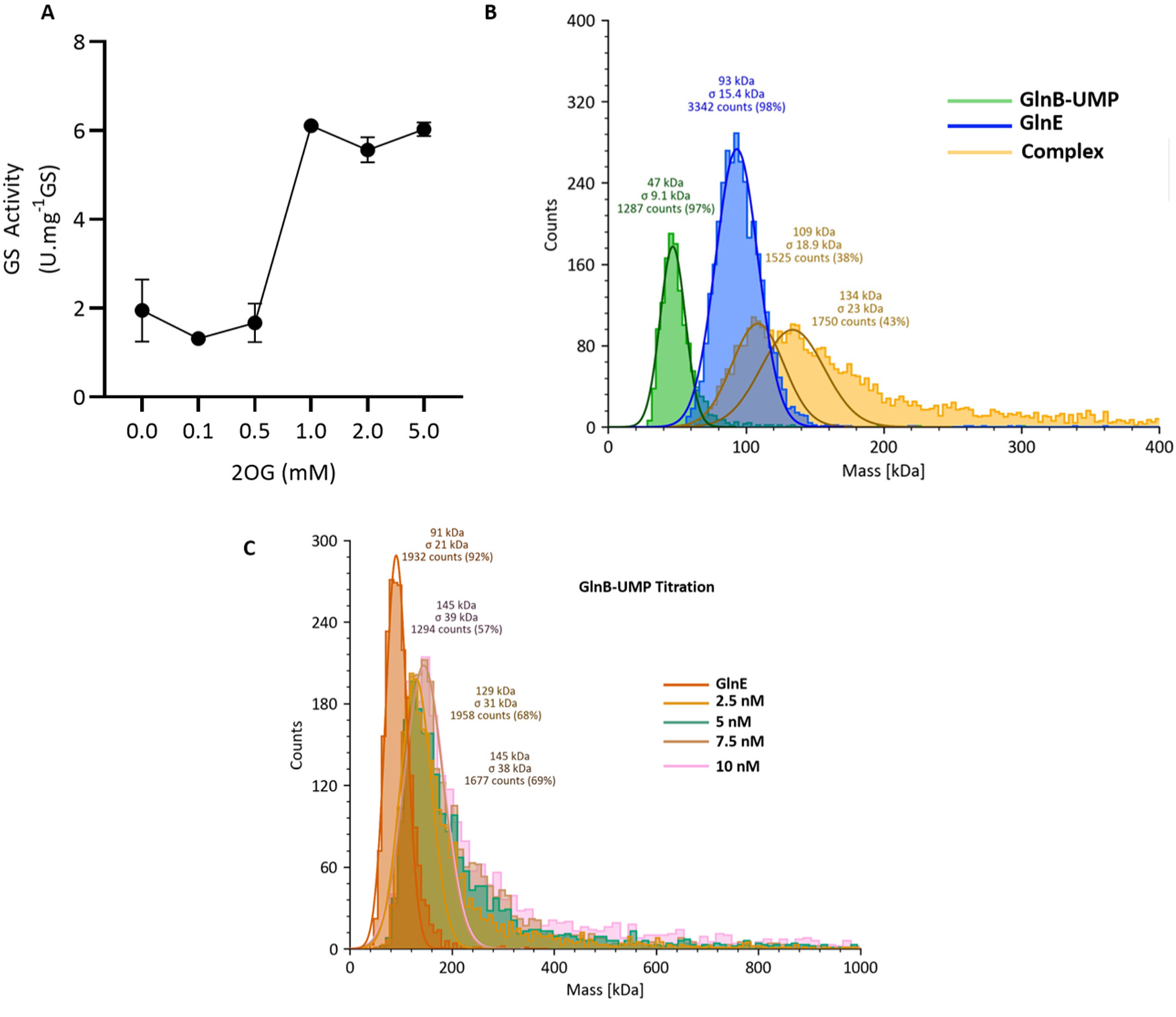
2OG modulation of GlnE: GlnB-UMP complex assembly. a) PK/LDH-coupled GS activity assay showed inhibition of GS activity up to 0.5 mM of 2OG. At concentrations higher than 1 mM, the AMPylation was inhibited, suggesting the GlnE: GlnB-UMP complex assembly. B) A mass shift observed in the mass photometry suggests a mass of 134 kDa for the GlnE:GlnB-UMP complex in the presence of 2OG. This experiment contained 10 nM of GlnB-UMP and GlnE, 5 mM of ATP, and 2OG. The same mass shift was observed for GlnB-UMP titration in the same condition (c). Taken together, the mass photometry indicates a 1:1 stoichiometry for the GlnE: GlnB-UMP complex. The GS activity was carried out in triplicate, and the results are described as the mean ± SD.

### The overall HsGS structure in adenylylated and non-adenylylated states

Despite the extensive work on GS, the structural mechanisms of how the AMPylation modulates its activity remains elusive. To solve this long-standing question, we used single particle cryo-EM to solve the structure of fully adenylylated HsGS (GS-AMP) in complex with either MgATP or MnADP and compared to unmodified GS in complex with MgATP (figure S6-S9). We solved the GS-AMP structure in complex with MnADP to investigate the mechanism of reverse γ-glutamyl-transferase (γ-GT) activity at the molecular level, whereas the structure of GS-AMP in complex with MgATP aimed at describing how AMPylation modulates GS activity under biosynthetic conditions. The overall resolution for GS+MgATP, GS-AMP+MgATP, and GS-AMP+MnADP were 1.71 Å, 1.77 Å, and 1.76 Å, respectively (supplementary table 1).

Similar to other type 1 GS enzymes, HsGS possesses a dodecameric assembly, composed of two hexameric rings (figures 5A, 5B, S10). The HsGS monomer consists of 471 amino acid residues structured as 15 β-strands and 20 α-helices, divided into an N-terminal the β-GRASP domain (residues 15-100) and a catalytic C-terminal domain (residues 107-471) (figure 5B). The β-GRASP domain is preceded by a solvent-exposed N-terminal helix (1-14) that projects outwards from the structure. As found in other GS enzymes, the catalytic domain harbors the bifunnel-shaped active site formed by the interaction of two adjacent monomers within the same hexameric ring. The active site is composed of Glu-loop (residues 331–336), Asn-loop (residues 258–270), and Tyr-loop (residues 156– 191) from each subunit, as well as the Asp-loop (residues 57–66) located in the β-GRASP domain of the adjacent monomer. The catalytic domain also contains the AMP-loop (387-404) and the C-terminal helix (461-471), the latter of which is embedded in a hydrophobic cavity of the opposing subunit from the other hexameric ring and participates in the dodecamer assembly.

**Figure 5.**
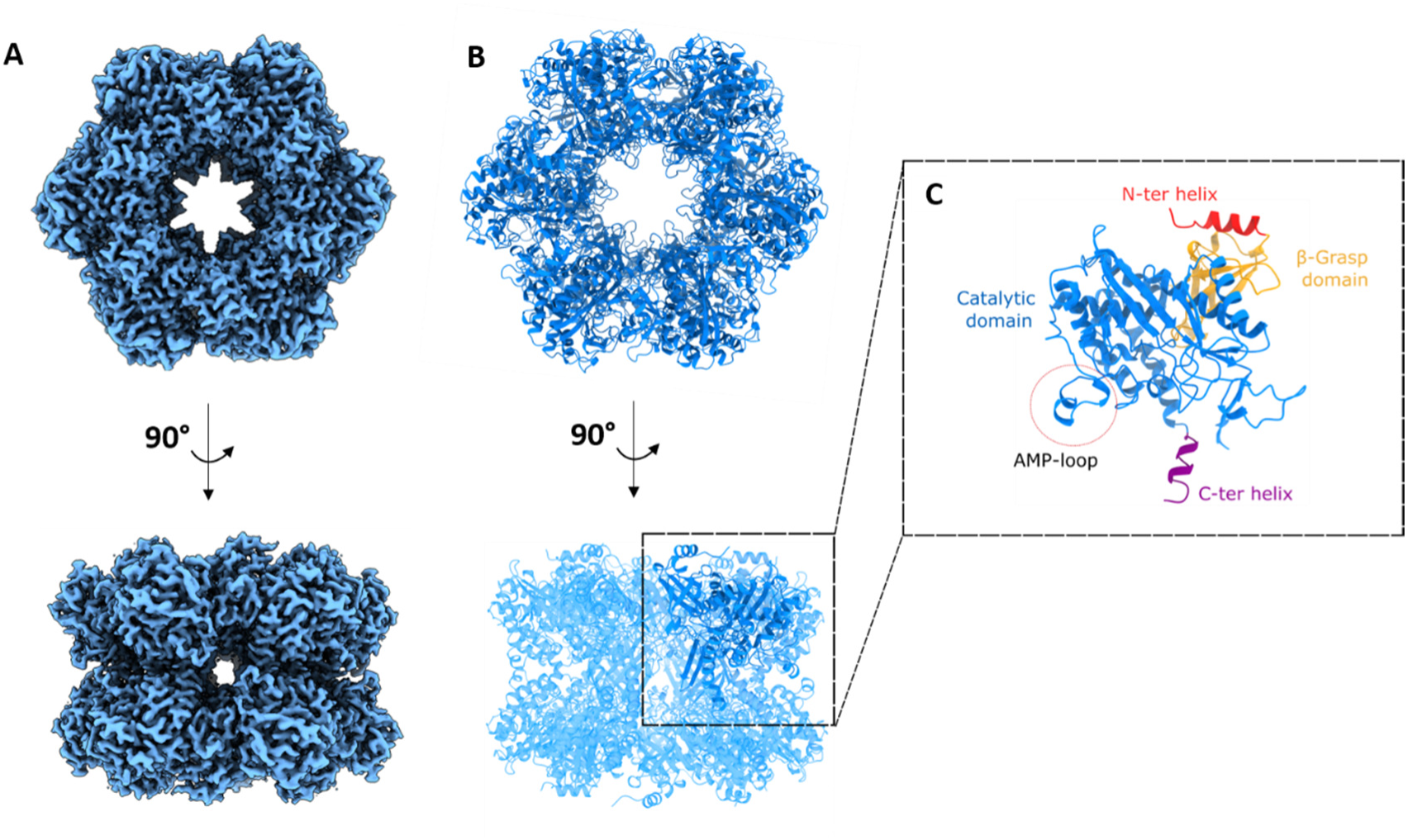
Overall GS structure of *H. seropedicae*. A) HsGS has a dodecameric assembly, characteristic of type 1 GS, as shown in the surface view. B) HsGS monomer is composed of the N-terminal helix (red), followed by the β-GRASP domain (yellow), which is connected to the catalytic domain (blue). In the catalytic domains is placed the AMP-loop, where the Y400 residue is target of AMPylation. The C-terminal helix (purple) connects is embedded in a hydrophobic cavity of the opposing subunit from the other hexameric ring and participates in the dodecamer assembly.

The comparison between the unmodified and adenylylated GS revealed that the overall structure of the enzyme was unaffected by AMPylation, with RMSD of 0.38 Å and 0.28 Å between unmodified GS and GS-AMP+MgATP or GS-AMP+MnATP, respectively (figure S11). However, while the AMP-loop was fully structured in the unmodified GS it was disordered in adenylylated GS complexes, as indicated by the absence of clear electron density in this region, suggesting that the addition of the AMP moiety increases the flexibility of the AMP-loop (figure 6A, S12-S14). Consequently, the sequence 396–408 (TKDL**Y**HLPPEEDK), which includes the AMPylation site Tyr400, could not be modeled. A second structural variation occurs within the segment 326–342 of adenylylated structures relative to unmodified GS (figure 6B). In unmodified GS, this segment is directly connected to the AMP-loop through two hydrogen bonds formed between Lys397-Asn341, and Leu399-Ser343 (figure 6C). Accordingly, the flexibility of this segment is likely a consequence of the disordered AMP-loop in both modified GS structures. While almost all residues in the 326–342 segment were resolved, the electron density for the Arg342 side chain was absent in adenylylated structures, indicating increased local flexibility of this residue upon adenylylation (figure 6D). Taken together, our data suggests that the AMPylation primarily induces flexibility in the AMP-loop, which subsequently alters the conformation of 326-342 segment via hydrogen-bod network and ultimately affects the positioning of the Arg342 side chain.

**Figure 6.**
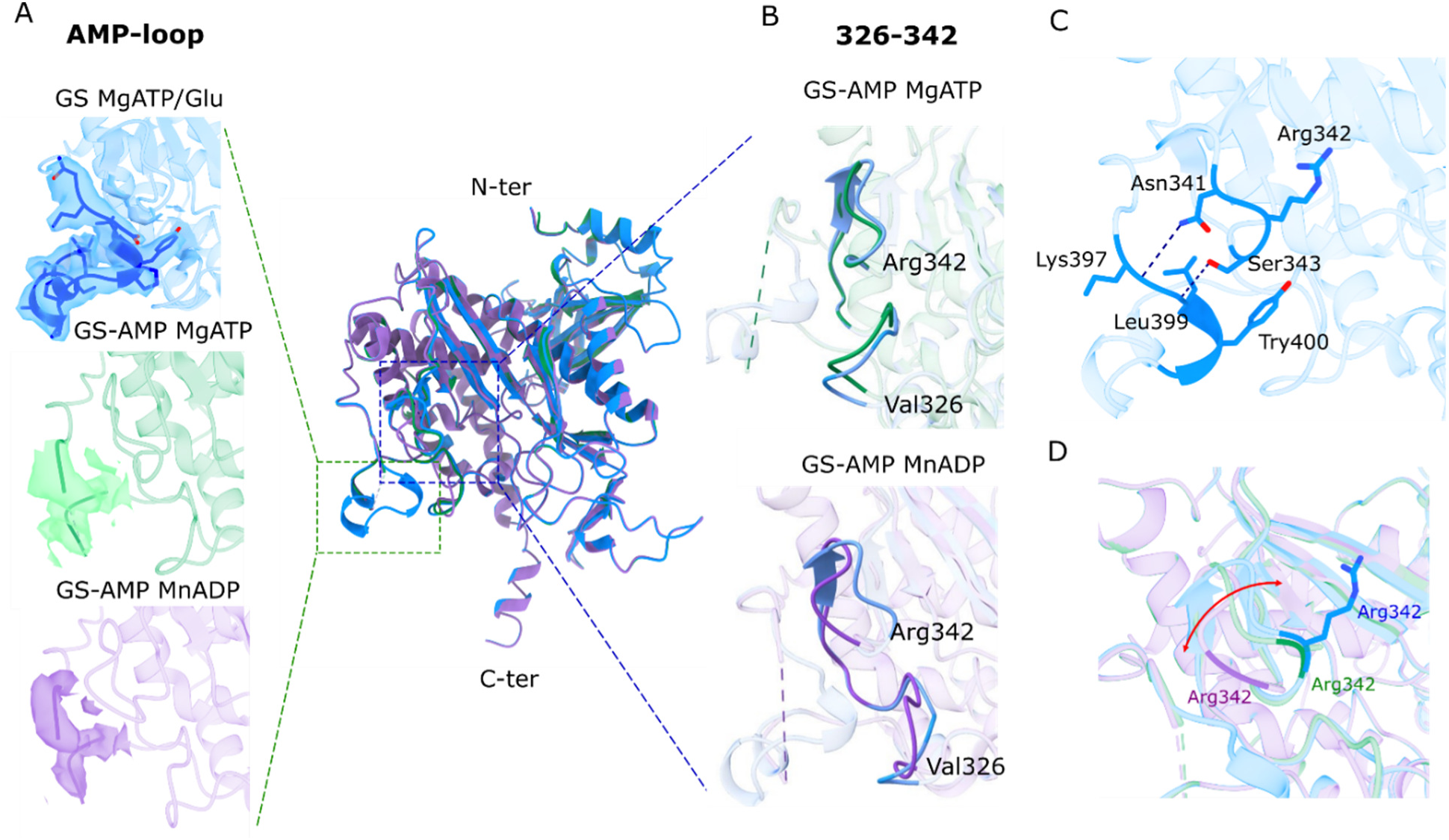
Structural comparison among GS monomers. A) GS AMPylation resulted in a loss of electron-density within the GS-AMP+MgATP (green) and GS-AMP+MnADP (purple), relative to unmodified GS (blue); consequently, residues 396–408 were not modeled in the adenylylated structures. B) Conformational changes induced by the AMPylation within the segment 326-342 in both GS-AMP+MgATP (green) and GS-AMP+MnADP (purple). C) In the unmodified GS, the AMP-loop and the segment 326-342 are stabilized by two hydrogen bonds between Lys397-Ans341 and Leu399-Ser343, linking the increased AMP-loop flexibility to the remodeling of 326-342 segment observed upon AMPylation. D) The movement of segment 326-342 increases Arg342 side-chain flexibility, as evidenced by the loss of electron-density in the adenylylated structures (in purple and green). The red arrow indicates the Arg342 movement among all three structures.

### Substrates binding into the HsGS and HsGS-AMP active sites

In the unmodified HsGS, the ATP is stabilized in the active site by the residues Phe228, Ser276, Arg358, Thr226, Glu210, Arg347, Arg362, and Arg342 (figure 7A and S15). Specifically, the adenine ring of ATP establishes three interactions: a hydrogen bond with Ser276 (3.55 Å); a π-π stacked interaction with Phe228 (within 4.4-4.5 Å); a cation-π interaction with Arg358 (3.90 Å). Additionally, Thr226 and Glu210 form hydrogen bonds with the 3’ (3.55 Å) and 2’ (3.06 Å) hydroxyl groups of the ribose moiety, respectively. The β and γ phosphates of ATP are coordinated with two Mg^2+^ ions that are separated by 5.74 Å from each other. The 1^st^ Mg^2+^ interacts with the γ (1.66 Å) and β (2.90 Å) phosphates and is coordinated by interactions with the residues Glu132 (2.77 Å), Glu360 (2.56 Å), and His272 (2.40 Å). Meanwhile, the 2^nd^ Mg^2+^ binds the γ phosphate (3.4 Å) and is coordinated by Glu134 (2.65 Å), Glu215 (2.68 Å), and Glu223 (2.42 Å). Importantly, the β phosphate is further stabilized by Arg347 through an electrostatic interaction (3.66 Å) and a hydrogen bond (3.48 Å), while Arg342 interacts with the γ phosphate via an electrostatic interaction at 3.19 Å (figure 7B).

**Figure 7.**
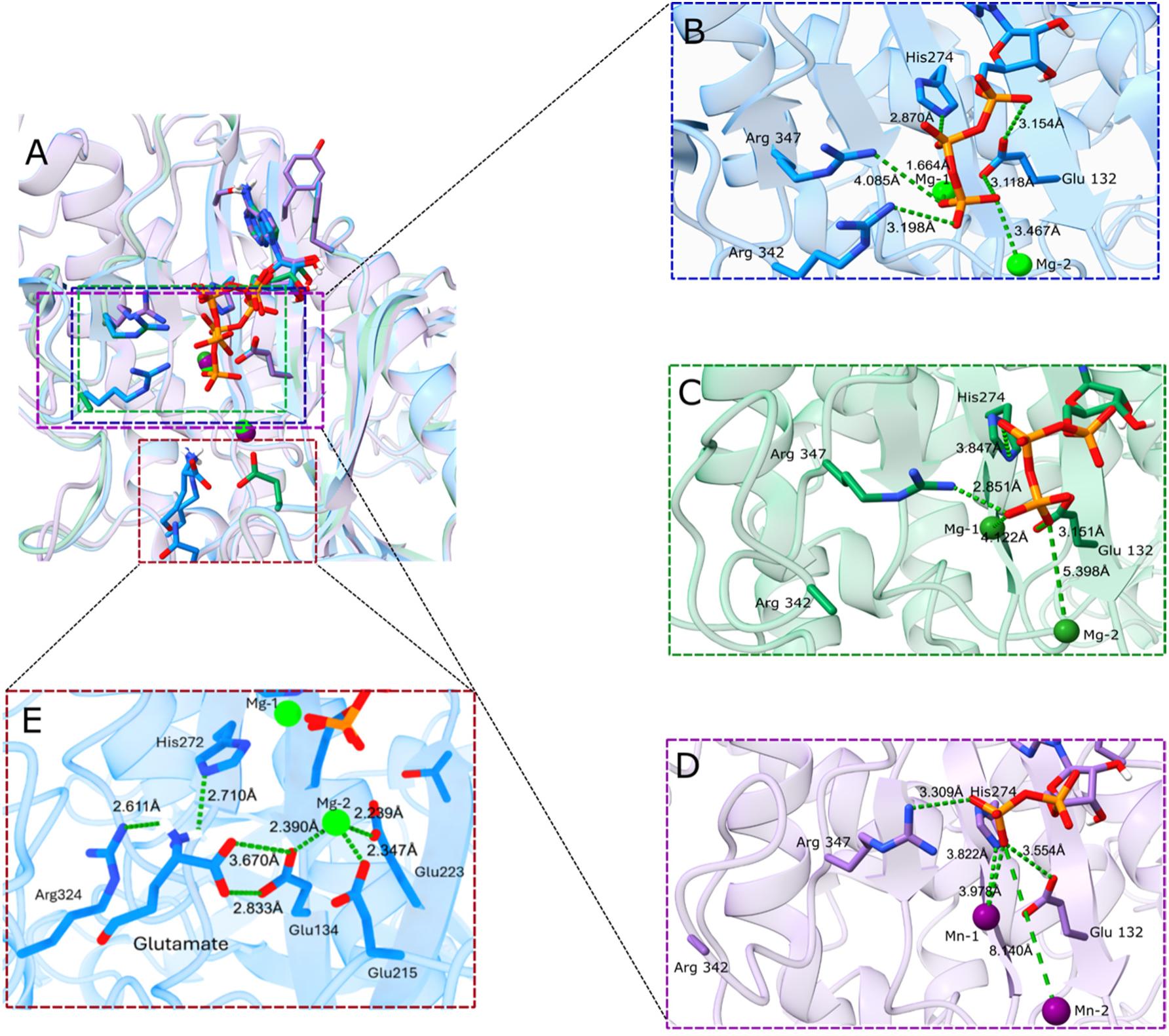
Nucleotide and glutamate binding sites in the GS and GS-AMP. A) Structural superimposition of all three structures indicates that while the nucleotides occupy identical binding pocket, their phosphate chains diverge significantly in position. B) In the unmodified GS, ATP adopts an extended conformation, with the γ-phosphate coordinated by 2^nd^ Mg^2+^ and directly interacting with Arg342. C) In the GS-AMP+MgATP structure, the ATP adopts a compacted conformation with the γ-phosphate distant from the Mg^2+^ and the interaction with Arg342 is disrupted. D) In the GS-AMP+MnADP structure, the β-phosphate of ADP does not interact with the 2^nd^ Mn^2+^ and the interaction with Arg342 is also disrupted. E) In the non-catalytic conformation, the main chain of glutamate binds to Glu134 and Arg324, but its side chain is far from the center of the active site, which would prevent the catalytic process from occurring.

The GS AMPylation significantly alters ATP and ADP binding within the active site. While the phosphate groups adopt an extended conformation in unmodified GS, they are arranged in a compact conformation in the adenylylated enzyme (figure 7C and D). In the GS-AMP+MgATP structure, the γ phosphate loses its interaction with the 2^nd^ Mg^2+^, with the ineratomic distance increasing from 3.46 Å to 6.07 Å. Similarly, in GS-AMP+MnADP structure, the 2^nd^ Mn^2+^ no longer coordinates the β phosphate, which are separated by 8.86 Å far from each other.

Another key difference is the position of Arg342 within the unmodified GS and GS-AMP structures. In the AMPylated structures, the side chain of Agr342 could not be resolved, indicating increased flexibility relative to unmodified GS. The disorder of Arg342 resulted in the loss of its interaction with the γ phosphate of ATP, increasing the distance to 11.72 Å (estimated from the main-chain position) and explaining the observed compact conformation of ATP. Likewise, the distances between Arg342 and β phosphate of ADP increased also to 10.47 Å.

Interestingly, we observed also a clear electron-density for glutamate only in the unmodified GS, occupying the lower part of the active bifunnel site. However, the orientation of its side chain suggests that the molecule was trapped in a non-productive conformation, located far from the Mg^2+^ in the catalytic center (figure 7E and figure S16). This observation indicates that the glutamate-binding site can accommodate the substrate in a different orientation distinct from that required for catalysis, pointing to a potential conformational rearrangement of the substrate before the reaction proceeding. In both GS-AMP structures, no electron density could be resolved for glutamate/glutamine, suggesting that GS AMPylation reduces substrate affinity despite identical sample preparation conditions to unmodified GS. The reduction in the affinity by the glutamate could be a caused by the differential nucleotide binding, as shown previously (Tomazini, et al. 2025).

## Discussion

Proteobacteria assimilate ammonium into glutamine and glutamate, with the GS– GOGAT cycle being particularly important under nitrogen limitation because GS has a much lower K_M_ for ammonium than GDH (Reitzer, 2003). Using a PK/LDH-coupled assay, we determined the apparent K_M_ of HsGS to be 0.15 mM for ammonium and 0.80 mM for glutamate in the presence of Mg²⁺ (Fig. 1A and 1B). The K_M_ for ammonium is comparable to values reported for other organisms, whereas that for glutamate appears lower than previously reported values (van Heeswijk et al., 2013).

The high affinity of GS for ammonium, together with the relatively high intracellular concentrations of glutamate and ATP, creates a potential metabolic cost under nitrogen-sufficient conditions. Glutamate is the most abundant amino acid in bacteria, reaching approximately 96 mM in *E. coli* growing under nitrogen-sufficient conditions (Bennett et al., 2009), although its intracellular pool is tightly buffered against metabolic fluctuations (Schumacher et al., 2013). Likewise, intracellular ATP concentrations can reach approximately 9.6 mM (Bennett et al., 2009). Thus, GS could potentially remain active and consume both ATP and glutamate when ammonium is abundant, necessitating tight regulation of the enzyme to prevent futile energy expenditure and perturbation of glutamate homeostasis.

In proteobacteria, GS regulation is not generally mediated by direct feedback inhibition by glutamine, but rather through a cumulative allosteric mechanism involving the end products of glutamine metabolism. In *H. seropedicae*, only tryptophan and threonine have been reported to inhibit unmodified GS (Tomazini et al., 2025). In addition to these allosteric mechanisms, ammonium availability triggers a rapid and reversible inactivation of GS *in vivo*, as widely reported in other organisms (Schutt and Holzer, 1972; Wax et al., 1982) and demonstrated here for *H. seropedicae* (Fig. 1C). GS activity decreased approximately three-fold within 30 sec after ammonium addition and was fully restored after 2 min. These observations prompted us to investigate the regulatory mechanism responsible for this rapid switch.

Our results indicate that, as in other bacteria, GlnE-mediated AMPylation contributes to the rapid regulation of HsGS. HsGS was progressively inactivated by increasing concentrations of GlnE *in vitro*, consistent with increasing AMPylation (Fig. 1D). Because the reaction mixture contained GS substrates (ammonium, ATP, and glutamate), this activity occurred in the absence of exogenous PII or glutamine. This observation could indicate that GlnE AT activity in *H. seropedicae* does not require exogenous glutamine under these conditions, although glutamine produced during the assay may also have contributed to GlnE activation.

Consistent with this possibility, exogenous glutamine alone did not affect GS activity, whereas glutamine progressively inhibited GS in the presence of truncated ΔNT-GlnE, corresponding to the C-terminal domain of GlnE (Fig. 2A and 2B). These results indicate that GlnE is responsive to glutamine in *H. seropedicae* and that the glutamine-responsive site resides within the C-terminal domain, as previously established for *E. coli*. Thus, although the general regulatory principle of glutamine-dependent GS AMPylation is conserved, our results define its operation in *H. seropedicae* and provide additional insight into the molecular signals controlling this process.

The DSF experiments further indicate that GlnE integrates glutamine and energy status through the interplay between MgATP and glutamine binding (Fig. 3). Glutamine alone produced little evidence of binding to ΔNT-GlnE, whereas MgATP bound with relatively low affinity. Pre-saturation with MgATP enabled glutamine binding, while pre-saturation with glutamine increased the apparent affinity for MgATP. These observations are consistent with cooperative or sequential binding of the two ligands, in which MgATP availability facilitates glutamine sensing and glutamine reciprocally enhances MgATP binding. Such coupling provides a potential mechanism by which GlnE activity can be constrained by cellular energy status. Because GS AMPylation is itself an ATP-dependent process, linking glutamine sensing to ATP availability could prevent unnecessary investment of energy in GS inactivation when cellular energy is limiting.

We next investigated the contribution of the PII protein GlnB to GlnE regulation. In *E. coli*, GlnB controls GlnE activity according to its UMPylation state (Atkinson and Ninfa, 1998; Jiang and Ninfa, 2009; Jiang et al., 2007), whereas GlnB-UMP has been reported to have little effect on GlnE in *R. rubrum* (Jonsson et al., 2007). In *H. seropedicae*, unmodified GlnB did not affect GlnE activity in the presence of ATP, ADP, or 2OG (Fig. 4A and 4B), consistent with previous evidence that GS AMPylation can occur independently of PII (Persuhn et al., 2000). Thus, the PII-dependent signal transduction between the N- and C-terminal domains of GlnE described in *E. coli* and *R. rubrum* is not essential for AT activity in *H. seropedicae*.

In contrast, our results demonstrate that the UMPylation state of GlnB provides an inhibitory signal to GlnE under conditions associated with nitrogen limitation. In *H. seropedicae*, GlnD promotes GlnB UMPylation under nitrogen-limiting conditions (Emori et al., 2018; Bonatto et al., 2007). Here, we show that HsGS AMPylation is inhibited by GlnB-UMP in the presence of ATP and 2OG (Fig. 4A and 4C). The inhibitory effect increased with 2OG concentration, and GS was not detectably modified at 1 mM 2OG, indicating formation of a GlnB:GlnE complex under these conditions (Fig. 4A). This concentration is physiologically relevant as the intracellular 2OG in *H. seropedicae* was reported to be approximately 4.5 mM before ammonium shock and to decrease to approximately 1.5 mM within 1 min after ammonium addition (Oliveira et al., 2015). Moreover, the reported K_D_ of 2OG for the third site of GlnB-UMP is 0.1686 mM (Oliveira et al., 2015), supporting the possibility that the GlnE complex forms in vivo.

The regulatory effects of GlnB-UMP are consistent with a model in which modification of the GlnB T-loop alters its interaction with GlnE. Attachment of the UMP moiety to the GlnB T-loop renders this region disordered (Palanca and Rubio, 2017). In *H. seropedicae*, the resulting interaction with GlnE may promote a transition toward a closed conformation in which the N- and C-terminal domains interact. Such an interaction could inhibit AT activity by interfering with glutamine binding or access to the active site, as proposed for *E. coli* GlnE (Jiang and Ninfa, 2009). Alternatively, the conformational transition could favor deAMPylation through the N-terminal domain. The latter possibility remains to be experimentally tested in *H. seropedicae*. Together, these findings indicate that the conserved GlnE/GlnB regulatory framework is maintained in *H. seropedicae*, while species-specific features also present in the control of GlnE activity. Having established how metabolic signals regulate GlnE-dependent GS AMPylation, we next asked how this regulatory modification is translated into loss of GS catalytic activity. To address this question, we solved three structures of HsGS: unmodified GS bound to MgATP, and AMPylated GS bound either to MgATP or to MnADP. Like other type 1 GS enzymes, HsGS forms a dodecamer consisting of two hexameric rings, with the active sites located between adjacent monomers (Fig. 6A and 6B).

AMPylation did not produce a major rearrangement of the overall GS structure, as evidenced by low RMSD values comparing the modified to the unmodified enzyme. However, the AMP-loop could not be resolved in the AMPylated enzyme, whereas its position was clearly defined in the unmodified GS (Fig. 7A). This observation suggests that AMPylation primarily affects the local structural dynamics of this loop rather than the global architecture of GS. Interestingly, a similar increase in flexibility has been observed for the T-loop of GlnB following UMPylation (Palanca and Rubio, 2017), suggesting that increased local disorder may represent a recurring structural consequence of nucleotide modification.

The structural data further provides a mechanism linking AMPylation to inhibition of GS catalysis. In the unmodified enzyme, Arg342 interacts with the γ-phosphate of ATP and contributes to maintaining ATP in an extended conformation, with the γ-phosphate positioned toward the center of the active-site funnel. This arrangement also favors coordination of the ATP phosphates with the second Mg²⁺ ion and is therefore expected to facilitate transfer of the γ-phosphate to glutamate during formation of the γ-glutamyl-phosphate intermediate (Joo et al., 2018). Upon AMPylation, increased flexibility of the AMP-loop is associated with displacement of Arg342, disrupting its interaction with the γ-phosphate and consequently the coordination of the second Mg²⁺ ion. ATP then adopts a more compact conformation in which the γ-phosphate is no longer optimally positioned for catalysis (Fig. 8). A similar compact nucleotide configuration was observed with MnADP (Fig. 8D).

These structural observations provide a mechanistic explanation for the long-thought inhibition of GS by AMPylation. Because glutamate binding to GS depends on prior ATP binding (Tomazini et al., 2025), the AMPylation-induced alteration in ATP conformation may also prevent productive glutamate binding. Consistent with this model, glutamate was observed only in the structure of unmodified GS. Thus, rather than causing a large-scale conformational rearrangement of the enzyme, AMPylation appears to inhibit GS through a localized increase in AMP-loop flexibility that propagates to the active site, alters ATP positioning, and compromises the structural requirements for catalysis.

Taken together, our results extend the current understanding of the conserved GlnE/GlnB regulatory system in *H. seropedicae* and provide a structural mechanism for the final step of this regulatory pathway. Under nitrogen-limiting conditions, high 2OG promotes the formation of the GlnE complex, thereby inhibiting GlnE AT activity and maintaining GS in its active, unmodified state to support ammonium assimilation. Conversely, under nitrogen-sufficient conditions, glutamine and MgATP promote GlnE-dependent GS AMPylation. Attachment of AMP to Tyr400 increases the flexibility of the AMP-loop, leading to repositioning of Arg342 and alteration of ATP conformation within the active site. This ultimately renders ATP incompetent for catalysis and inhibits GS activity. Thus, the present findings connect the established metabolic regulation of GS AMPylation to its previously unresolved structural consequences, providing a mechanistic link between cellular nitrogen sensing and the catalytic state of GS in *H. seropedicae*.

## Material and Method

### GS switch-off

*H. seropedicae* SmR1 was cultivated in NFbHP-Malate medium (Klassen, G., et al. 1997) supplemented with 5 mM glutamate for 24 hours at 30°C, 150 rpm. Immediately before the switch-off induced by ammonium shock, a 20 μL aliquot was taken (time 0). After that, 200 μM NH₄Cl was added. Subsequent 20 μL aliquots were collected at 0.5, 1, 2, 5, and 30 minutes. These aliquots were flash-frozen in liquid nitrogen. After freezing, the samples were centrifuged, and the cells were resuspended in 400 μL of 1% KCl and lysed by sonication on ice for 2 minutes. The cell lysates from all-time points were used to determine γ-glutamyl-transferase GS activity as described below.

### Cloning

The *glnE* gene encoding the GlnE enzyme was PCR-amplified from the *H. seropedicae* SmR1 genome using the following primers: *glnE_fwd* 5’-CATCGACATATGGCCGCCGGTTTTCCTTC-3’; *glnE_rev*5’-GTCAGGATCCTCAGCCAAAGACTTGCTGCCA-3’. To delete of the N-terminal domain (adenylyl-removing domain) of GlnE to create the unidirectional adenylyl-transferase ΔNT-GlnE, the fragment of the glnE gene was PCR-amplified using the following primers: *glnE3_fwd* 5’-CATCGACATCATATGGCCGACAAGCAGTCCGA-3’; *glnE_rev* 5’-GTCAGGATCCTCAGCCAAAGACTTGCTGCCA-3’. Both *glnE* and *ΔNT-GlnE* PCR products were digested with NdeI and BamHI restriction enzymes and ligated to the pETNdeM-11 (GlnE) (Little, R. et al., 2011) or pET28a (ΔNT-GlnE) vectors previously digested with NdeI/BamHI, yielding the plasmids *glnE*pETNdeM11 and *ΔNT-GlnE*pET28a expressing N-terminal 6x His tag version of GlnE and ΔNT-GlnE, respectively.

### Protein expression

The plasmids *glnA*pETNdeM-11 (HsGS) (Tomazini, L.F. et al, 2025), *glnE*pETNdeM-11, *ΔNT-GlnE*pET28, and pETHsglnB (HsGlnB) (Stefanello, A.A., et al., 2020) were transformed into chemically competent *E. coli* BL21(DE3) cells by chemical transformation (Sambrook, J. et al., 1989). Unmodified GS was expressed in glutamine-supplemented M9 medium as previously described (Tomazini, L.F. et al, 2025). The expression of GlnB was carried out in LB medium supplemented with 20 mM NH_4_Cl as previously described (Oliveira, M.A.S. et al, 2015). Briefly, both GS and GlnB grew under shaking at 150 rpm and 37°C until OD_600_ 0.4. Then, the expression started by adding 0.5 mM IPTG and took place under shaking at 150 rpm and 16°C for 3 h. The cells were harvested by centrifugation at 10.800 *g* and 4°C for 10 min.

For expression of GlnE and ΔNT-GlnE, 500 μL of an overnight-grown pre-culture was transferred to 500 mL of LB medium (Sambrook, J. et al, 1989) and kept shaking at 150 rpm and 37°C until OD_600_ 0.4. The cultures were then kept in an ice bath for 30 min, and the expressions were started by adding 0.5 mM IPTG. Expression took place under shaking at 150 rpm and 16°C for 16 h. The cells were harvested by centrifugation at 10.800 *g* and 4°C for 10 min.

### Protein purification

Cell pellets containing the overexpressed proteins were resuspended in buffer A (50 mM TRIS, pH 8; 50 mM NaCl; 20 mM Imidazole) and kept in an ice bath with 0.5 mM PMSF for 15 min. Cells were lysed by sonication in an ice bath for 5 min, with 5 s pulses and 5 s intervals. Alternatively, cell lysis could also be performed using a French press. The soluble and insoluble fractions were separated by centrifugation at 10.800 *g* for 10 min at 4°C.

The GS, GlnE, and ΔNT-GlnE proteins were purified by affinity chromatography. The soluble fraction was loaded onto a HiTrap-Chelating-Ni^2+^ column (Cytiva). The proteins were eluted using a gradient of 20 to 1000 mM imidazole in buffer B (50 mM TRIS, pH 8; 50 mM NaCl; 1 M Imidazole). The GlnE enzyme was submitted to Size Exclusion Chromatography in the Superdex 200 Increase 10/300 (Cytiva) column eluted with buffer 50 mM TRIS pH 8 and NaCl 150 mM.

For GlnB purification, the lysed extract was pre-heated at 70°C for 20 min, followed by centrifugation at 10.800 *g* for 10 min at 4°C. The soluble fraction was loaded onto a previously equilibrated HiTrap-Heparin chromatographic column (Cytiva). The proteins were eluted using a gradient of 20 to 1M NaCl in buffer 50 mM TRIS, pH 8.

In all cases, the fractions containing purified proteins were dialysed in two steps: first against a buffer containing 50 mM TRIS pH 8.0 and 50 mM NaCl; subsequently, the proteins were transferred to a buffer containing 50 mM TRIS pH 8.0, 50 mM NaCl, and 50% Glycerol for at least 12 hours. The proteins were quantified by spectrophotometry at 280 nm.

### GlnB uridylylation *in vitro*

*In vitro* uridylylation of GlnB was performed as previously described (Oliveira, M.A.S. et al., 2015) with modifications. For the uridylylation of 200 μM GlnB, the uridylylation mix was supplemented with 66 μL of Mg5x buffer (500 mM TRIS, pH 7.5; 500 mM KCl; 125 mM MnCl_2_), 5 mM 2OG, 1 mM UTP, 0.2 mM ATP, and 1 μM of previously purified GlnD enzyme. The reaction was incubated overnight at room temperature. The progress of uridylylation was monitored by non-denaturing polyacrylamide gel electrophoresis. At the end of the process, GlnD was inactivated by heating at 70°C for 15 min, followed by centrifugation at 10.800 *g* for 10 min at 4°C to separate GlnD and GlnB-UMP. Subsequently, the GlnB-UMP protein was dialyzed in a buffer containing 50 mM TRIS, pH 8.0, 50 mM NaCl, and 50% Glycerol for 12 hours.

### PK/LDH coupled GS assay

The GS activity was monitored using a system coupled to NADH consumption by the pyruvate kinase (PK) and lactate dehydrogenase (LDH) enzymes. For this, 50 nM of GS was mixed with the reaction mix (5 U PK; 5 U LDH; 20 mM MgCl_2_; 50 mM KCl; 1 mM phosphoenolpyruvate; 0.4 mM NADH; 50 mM Imidazole, pH 7; 0.5 mM NH_4_Cl; and 5 mM ATP). NADH consumption was monitored by spectrophotometry at 340 nm. The GS activity was expressed as U.mg^-1^GS. The experiments were carried out in triplicate, and the results are described as the mean ± SD.

### Biosynthetic GS assay

The post-translational state of GS was determined measuring both the biosynthetic and the γ-glutamyl-transferase (γ-GT) colorimetric activities as described previously (Tomazini, L.F. et al., 2025).

### *In vitro* GS AMPylation

The GS *in vitro* AMPylation was carried out in both PK/LDH and γ-GT assays. In the PK/LDH assay, in the reactional mix, GlnE or ΔNT-GlnE was added at a final concentration of 100 nM; GlnB or GlnB-UMP was used at 500 nM. When necessary, 2OG was added at 3 mM. The GS reactions were initiated by adding 0.5 mM glutamate after 15 min of AMPylation. NADH consumption was monitored by spectrophotometry at 340 nm.

In the γ-GT assay, 5 μL of the AMPylation mix (5 mM TRIS pH 7.5; 25 mM KH_2_PO_4_; 1 mM Glutamine; and 1 mM ATP) was added to unmodified GS (300 nM) and 100 nM of GlnE or ΔNT-GlnE. As required by the experiment, 200 nM of GlnB or GlnB-UMP were added. ATP, ADP, and 2OG were added at 5 mM final concentration as needed. The reaction proceeded at 37°C for 20 min. After this period, we proceeded with the γ-GT assay.

### Mass Photometry (MP)

MP measurements were carried out using the TwoMP mass photometer (Refeyn), as previously described (Samir et al., 2025b; Wellner, T. et al, 2026). Shortly, the instrument was calibrated with a 40 nM BSA/TG solution, diluted in interaction buffer (50 mM Hepes pH 7.5; 20 mM MgCl_2_; 300 mM NaCl), which was also used for focusing. To assess the interaction conditions, both GlnE and GlnB/GlnB-UMP proteins were pre-incubated at room temperature with the modulators ATP, ADP, or ATP/2OG in the interaction buffer. The final concentration of GlnE and GlnB/GlnB-UMP used was 10 nM, unless otherwise specified. For ATP, ADP, and ATP/2OG, the final concentration was 5 mM. Data was collected for 1 minute using AcquireMP software. The results were analyzed, and graphs were generated using DiscoverMP software, as shown previously in (Müller et al., 2026; Wellner, T. et al, 2026).

### Differential Scanning Fluorimetry (DSF)

Differential Scanning Fluorimetry (DSF) was used as described previously (Samir et al., 2025a) to characterize the interaction of allosteric regulators with the truncated GlnE 3-6 protein. Each protein-ligand interaction reaction contained 30 μM of protein, 9 µL of 5x buffer (500 mM Hepes; 500 mM NaCl; 125 mM MgCl₂), 5 mM SYPRO Orange (ThermoFisher), and enough water to reach a final volume of 20 µL. Ligands were titrated as indicated in the experiment. The reactions were carried out in a StepOne RT-PCR thermocycler (Applied Biosystems) over a temperature range of 25 to 94°C, with a 1°C increase per minute. Fluorescence emission spectra were generated using the StepOne v2.3 software and expressed as the reciprocal of the first derivative of fluorescence with respect to temperature (–dF/dT). The resulting peaks represent melting temperatures (Tm). After obtaining the melting temperatures, the variation in Tm (ΔTm) between the ligand condition and the ligand-free control was calculated (Nissen, F.H. et al.,2007).

### Cryo-EM

The general procedures for data acquisition and processing were done as described previously (Elshereef et al., 2026; Berwanger et al., 2026). In summary, the structures of unmodified GS in complex with MgATP and adenylylated GS in complex with MgATP or MnADP from *H. seropedicae* were solved through Cryo-EM. For that, unmodified GS were expressed and purified. To ensure a high degree of adenylylation, GS was adenylylated *in vitro* with a unidirectional adenylyl-transferase ΔNT-GlnE enzyme. For GS adenylylation, 2 mg of GS was mixed with AMPylation buffer (50 mM Tris, pH 7.5; 20 mM MgCl_2_; 1 mM Glutamine; 1 mM ATP; 72 μM ΔNT-GlnE) and incubated at 37°C for 1 h. The GS modification state was confirmed through the PK/LDH assay immediately before freezing, showing a complete inhibition of GS. Three microliters of 1.5 mg/mL purified protein were pipetted onto a glow-discharged Quantifoil R 1.2/1.3 300-mesh gold holey carbon grid (Quantifoil). Grids were blotted for 6 s with a blotting force of 5 and a humidity of 100% at 4 °C, and then flash-frozen into liquid nitrogen-cooled liquid ethane using a Mark IV Vitrobot (FEI). To avoid preferred orientation bias, 4 mM CHAPSO was added to prevent water-air interface interactions; accordingly, the protein concentration was increased to 15 mg/mL. The samples conteined TRIS 50 mM pH 7.5.

For unmodified GS, MgCl_2_, Glutamate, ATP and NH_4_Cl were added. For AMPylated GS, we prepared two samples: one with MgCl_2_, Glutamate, ATP and NH_4_Cl and other with MnCl_2_, Glutamine, ADP. In all cases, the final concentration of substrates was 5 mM. A 5 mM ATP was only added before freezing. Micrographs were acquired in EER-format on the Thermo Fisher Titan Krios G4 microscope equipped with a cold field-emission gun, a Selectris X energy filter, and a Falcon IVi detector set to a nominal magnification of 215,000X, corresponding to a pixel size of 0.572 Å operating in counting mode. The energy filter was operated with a slit width of 10 eV to remove inelastically scattered electrons. The total exposure time was 10 s, corresponding to a total dose of 40 e–/Å2. Forty individual frames were collected with an electron dose of 1.0 e–/Å2 per frame. Data was acquired with aberration-free image shift (AFIS) using EPU software at a throughput of 750 images per hour. Defocus range was −0.5 to −2.0 μm.

### Cryo-EM image processing

Cryo-EM processing was achieved as described previously (Haffner et al., 2023; Berwanger et al., 2026). In summary, a total of 13848 movies were collected on the Titan Krios G4 microscope equipped with a Falcon IVi detector. The movies were important for the cryoSPARC and were motion corrected using patch motion correction, followed by CTF estimation using the patch CTF estimation in cryoSPARC. Using blob picker, 2,795,951 particles were picked, subjected to several rounds of reference-free 2D classification, and particles included in the classes showing good alignment were subjected to *Ab-Initio* reconstruction. All the *ab initio* reconstruction volumes were used for Hetero Refinement. A class with a distinct volume was chosen, and particles belonging to that class were extracted at a rate of 1.27 Å/pix. After homogeneous 3D refinement in C1 symmetry, using the re-extracted particles, a 2.77 Å map was produced. By applying the D6 symmetry in NU-refinement using the re-extracted particles, a 2.34 Å map was produced. A second round of homogeneous refinement was conducted by applying D6 symmetry to obtain a 1.7 Å map. A soft mask was applied to the GS glutamine synthetase for local refinement to further improve the quality of density. Local resolution estimation is also performed in cryoSPARC. The structure was modelled using the final 1.7 Å map. Data processing for modified GS-AMP+MgATP and GS-AMP+MnADP was done similarly as that for the unmodified GS-ATP.

### Model building, refinement, and validation

AlphaFold was used to create an initial model using the protein sequences (Jumper et al., 2021), and UCSF Chimera was then used to fit the rigid bodies into the density (Pettersen et al., 2021). Coot was used to manually rebuild the model (Emsley et al., 2010). PHENIX was used to refine the final model in real space (Liebschner et al., 2019). UCSF ChimeraX was used to create model illustrations (Pettersen et al., 2021). All the figures were generated in ChimeraX and PyMOL. For the models of modified GS-AMP+MgATP and GS-AMP+MnADP, the atomic model of GS-ATP was fitted into the 3D density maps using UCSF Chimera and applied to refinement using phenix.real_space_refine and Coot as mentioned above. The statistics for the geometries of models were generated using MolProbity and summarized in the supplementary information Table S1.

## Data availability

Cryo-EM maps have been deposited in the wwPDB OneDep System under the Electron Microscopy Data Bank accession codes EMD-58843, EMD-58891 and EMD-58881 for the unmodified GS at 1.7 Å, modified GS in complex with MgATP at 1.77 Å, and modified GS in complex with MnADP 1.76 Å maps, respectively. The atomic models associated with the corresponding maps have been deposited in the PDB under accession codes 32EQ, 32HH and 32GU, respectfully.

## Acknowledgments

This work was funded by grants from the German Research Foundation (DFG) as part of the collaborative research centers SFB1381 (project number: 403222702; subproject B12) and Emmy Noether program (SE 3449/3-1) to KAS. We would like to thank the TEM and Cryo-EM facilities at the University of Freiburg, for microscope access and data collection. The TEM (Hitachi HT7800 at the Faculty of Biology) and the Titan Krios G4 cryo-TEM were funded by the DFG (project numbers 426849454 and 506518771, respectively) and are operated as a partner unit within the Life Imaging Center (LIC) and the Microscopy and Image Analysis Platform (MIAP). Also, we thank Daniel Wohlwend (Freiburg University) for facilitating the usage of mass photometry. This study was financed in part by the Coordenação de Aperfeiçoamento de Pessoal de Nível Superior - Brasil (CAPES) - Finance Code 001.

## Supplementary figures

**Figure S1.**
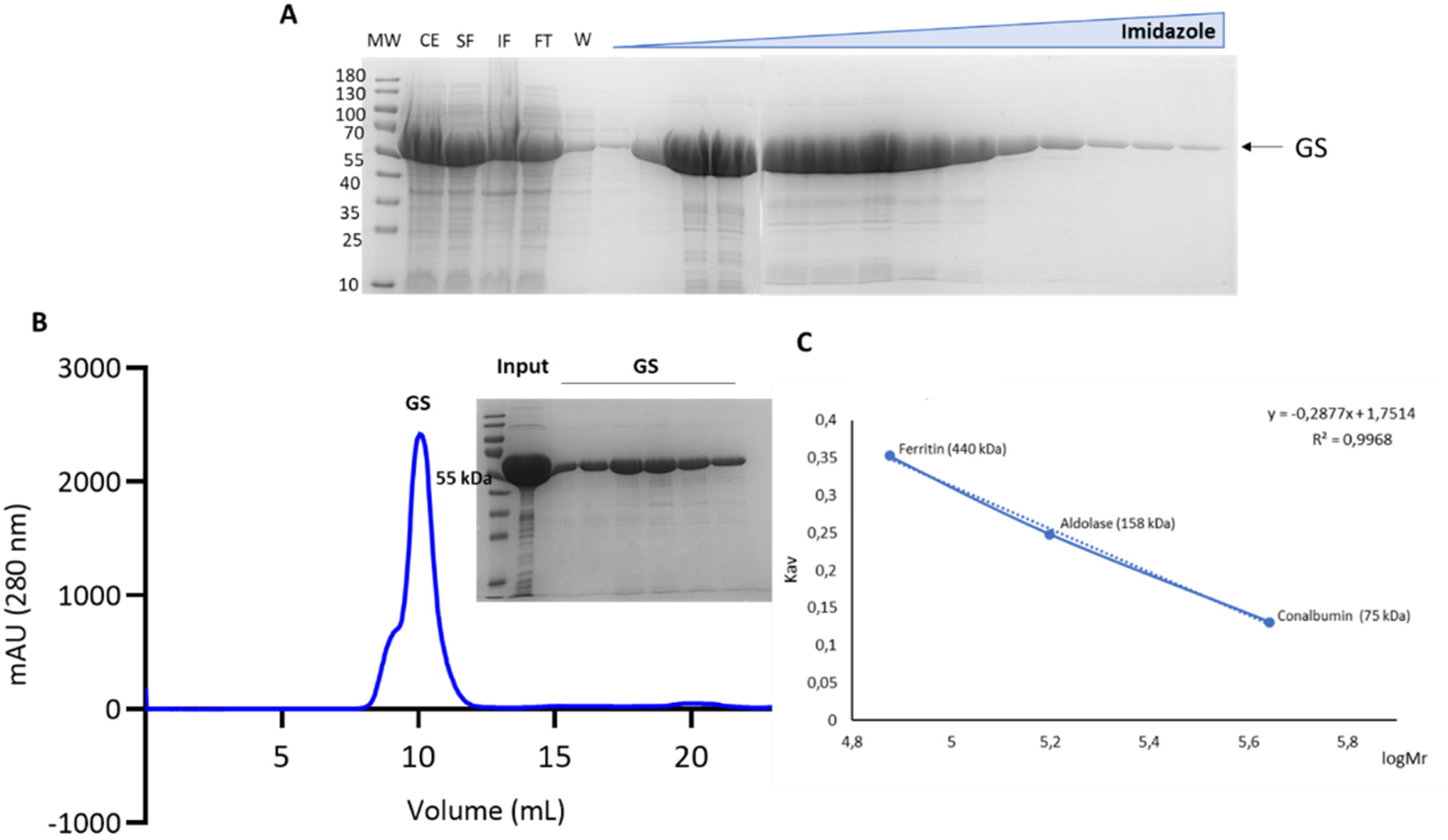
Glutamine synthetase purification. A) SDS-PAGE of GS purification by IMAC (Immobilized Metal Affinity Chromatography). The protein was eluted using an imidazole gradient. The final concentration of 14 mg/mL was determined spectrophotometry. MW: Molecular Weight Marker; CE: Crude Extract; SF: Soluble Fraction; IF: Insoluble Fraction; FT: Flow-through; W: Wash. B) After purification by IMAC, the GS was submitted to SEC, eluting at 10 mL. C) Calibration curve of the SEC used to estimate the molecular mass of GS as 597 kDa, corresponding likely to dodecamer.

**Figure S2.**
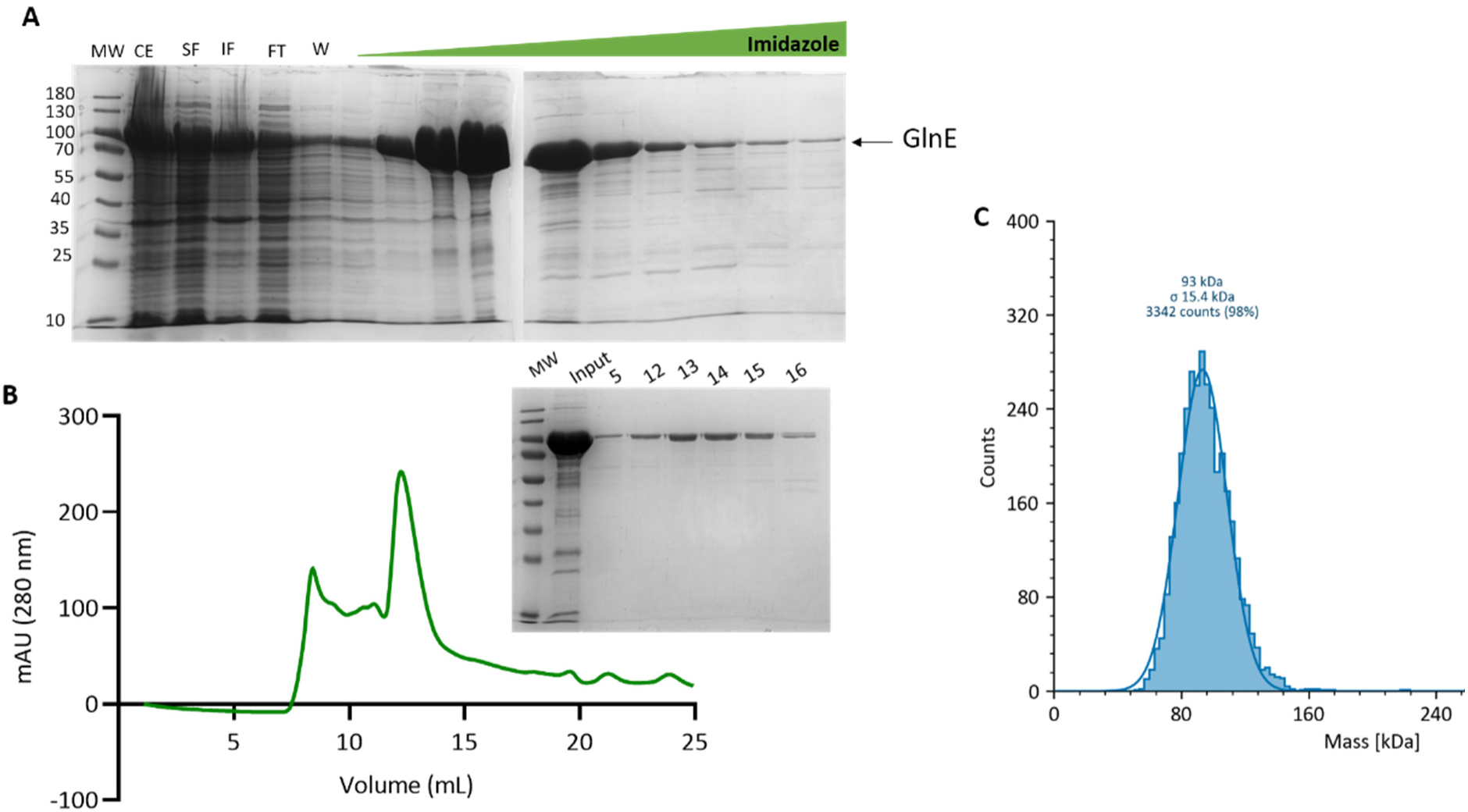
GlnE purification. A) SDS-PAGE of GlnE purification by IMAC (Immobilized Metal Affinity Chromatography). The protein was eluted using an imidazole gradient, with the main elution occurring between 216 mM and 416 mM. The final concentration of 4.9 mg/mL determined spectrophotometry. MW: Molecular Weight Marker; CE: Crude Extract; SF: Soluble Fraction; IF: Insoluble Fraction; FT: Flow-through; W: Wash. B) After purification by IMAC, GlnE was submitted to SEC, which suggested two populations. The first peak (around 7 mL) corresponds to aggregation, while the second peak (around 13 mL) corresponds to the GlnE monomer. C) The GlnE molecular mass of 93 ± 15 kDa (corresponding to a monomer) was further determined by mass photometry that employs physiological nM concentrations.

**Figure S3.**
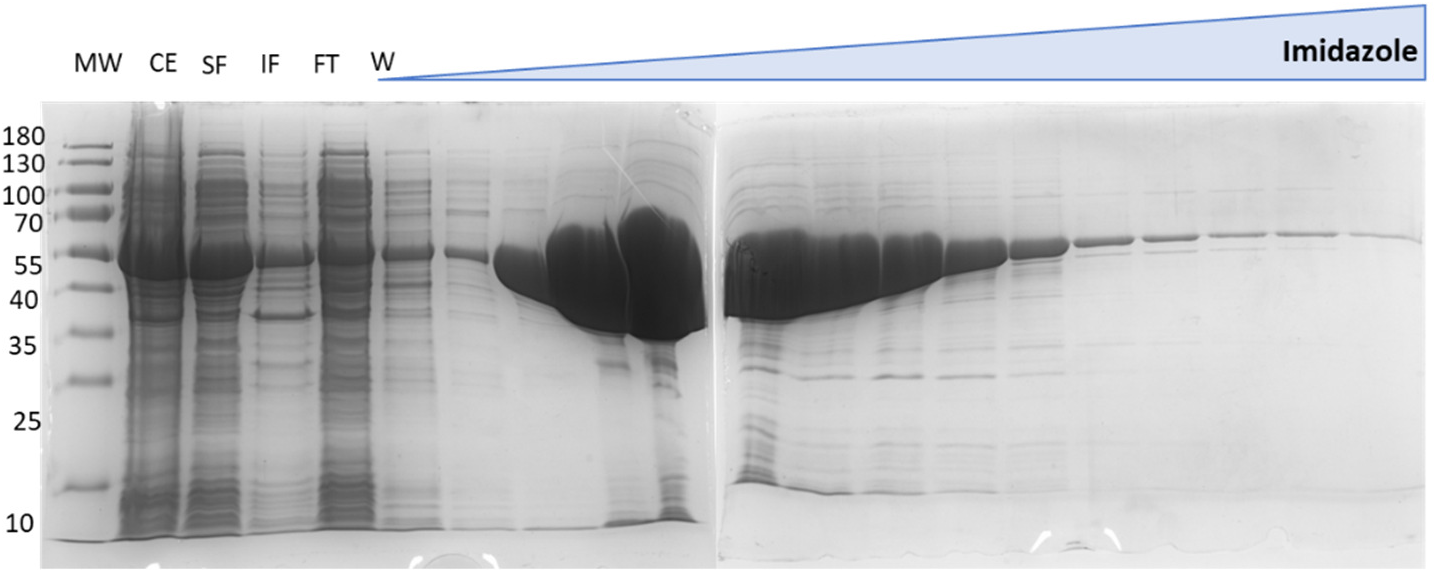
Purification of the *H. seropedicae* ΔNT-GlnE enzyme. SDS-PAGE of the affinity chromatography for ΔNT-GlnE purification. The protein eluted between 216 and 412 mM imidazole, as indicated by the bands at ∼55 kDa. MW: molecular weight marker; CE: crude extract; SF: soluble fraction; IF: insoluble fraction; FT: flow-through; W: wash.

**Figure S4.**
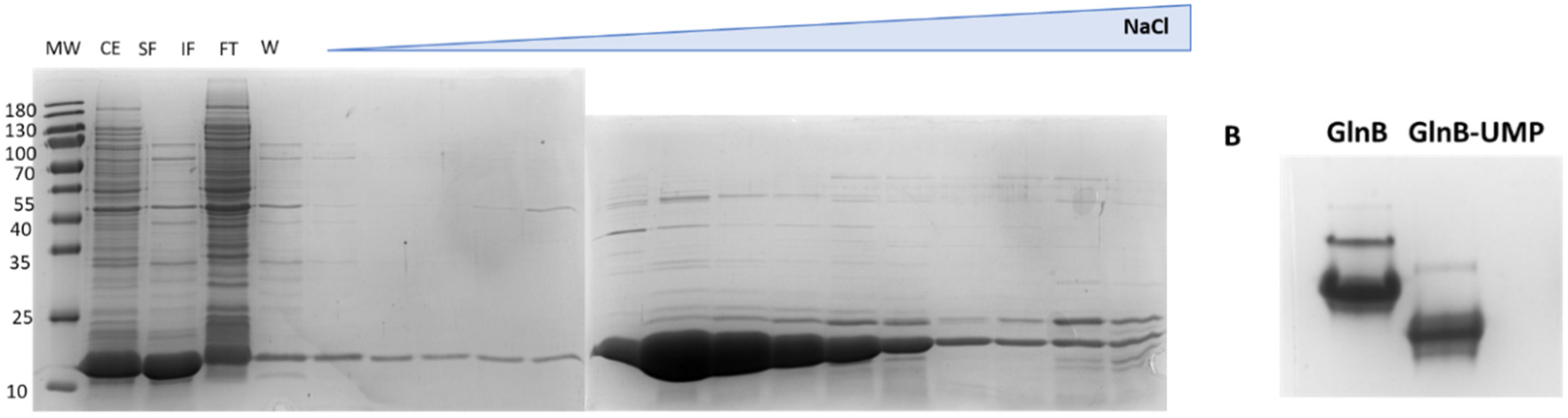
Purification and *in vitro* uridylation of the *H. seropedicae* GlnB protein. A) SDS-PAGE of the ion-exchange chromatography for GlnB purification. The bands at ∼13 kDa indicate that the protein was eluted between 421 and 608 mM NaCl. MW: molecular weight marker; CE: crude extract; SF: soluble fraction; IF: insoluble fraction; FT: flow-through; W: wash. B) Native-PAGE to verify the *in vitro* uridylation of GlnB. The shift in the electrophoretic profile indicates uridylation.

**Figure S5.**
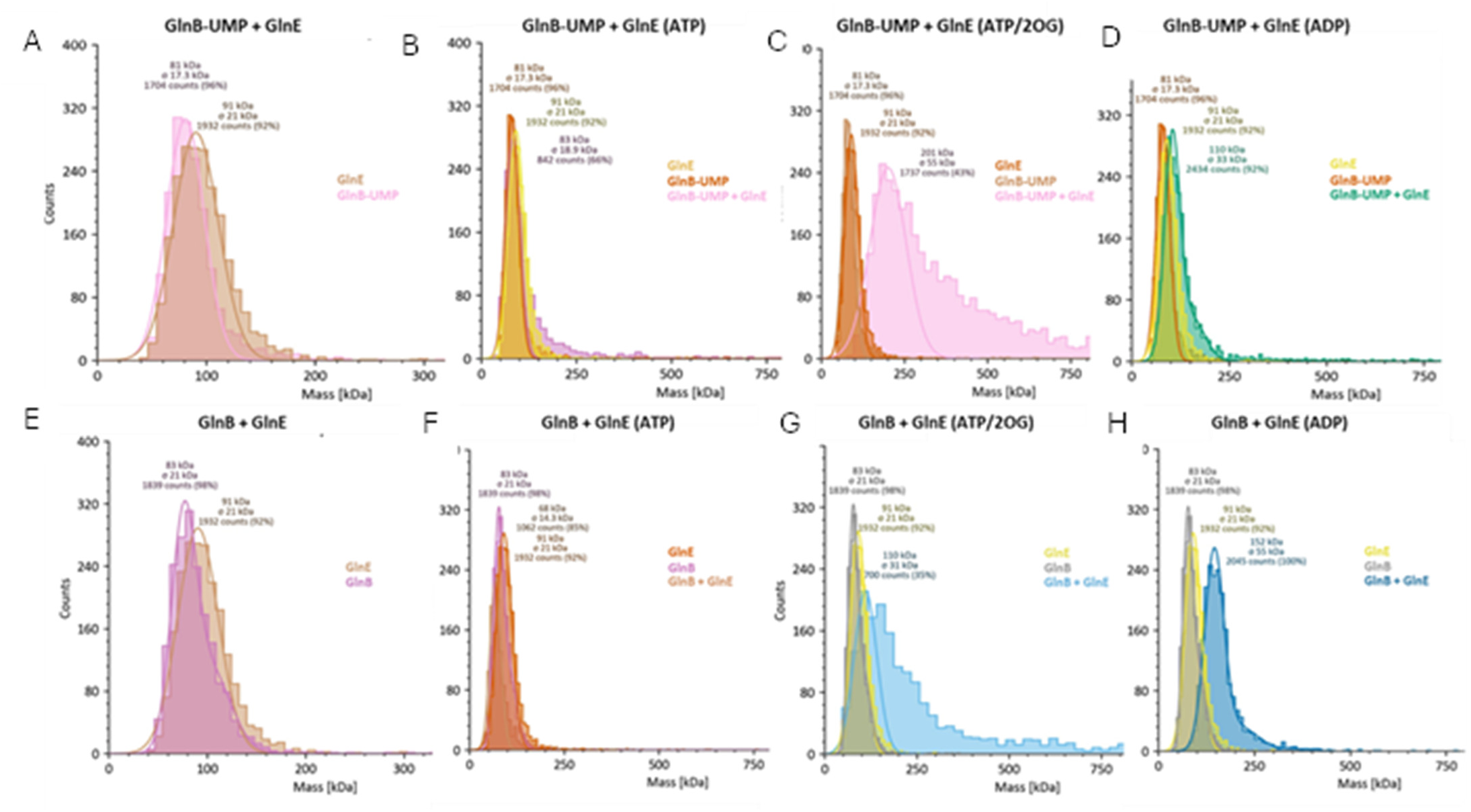
Interaction conditions for GlnE and GlnB/GlnB-UMP. All interaction conditions were tested independently. The mass shift under the GlnB-UMP+ATP+2OG condition was reproduced, suggesting complex formation. A and E) Overlap of the masses of the proteins used in each assay; B, C, and D) Conditions tested for the formation of the GlnB-UMP:GlnE complex; F, G, and H) Conditions tested for the formation of the GlnB:GlnE complex. Proteins were used at 10 nM (GlnE) and 20 nM (GlnB/GlnB-UMP). Effectors were added at 5 mM.

**Figure S6.**
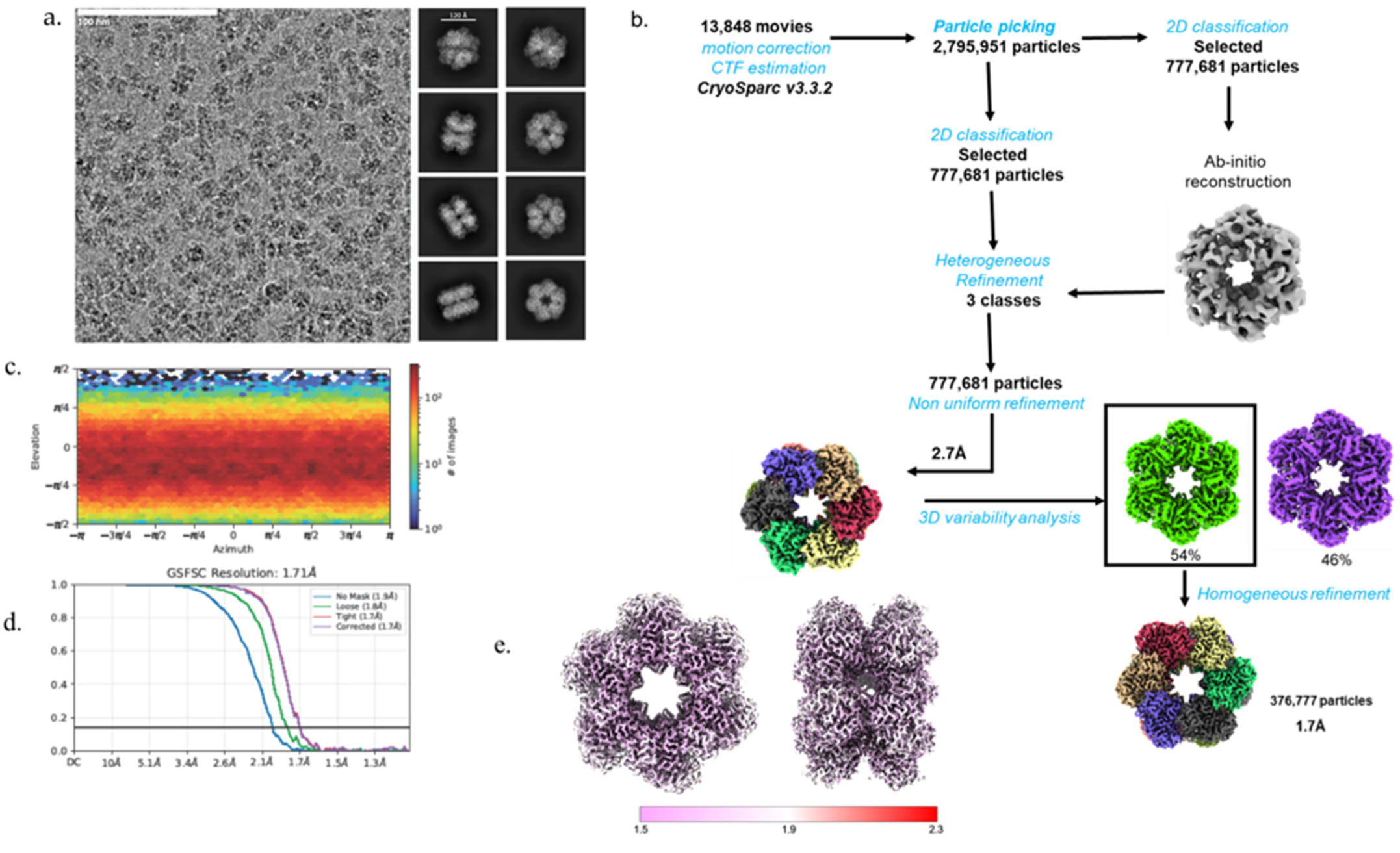
Processing workflow for unmodified GS in complex with MgATP and Glutamate. A) Representative motion-corrected micrograph showing different orientations of the GS+MgATP/Glu particles and 2D class averages; B) Cryo-EM processing workflow used for obtaining the high-resolution structure of GS+MgATP/Glu. C) Histogram showing the 2D distribution of the particle viewing directions used in the 3D reconstruction: D) Fourier shell correlation (FSC) curves. E) Local resolution maps for different orientations of GS+ATP/Glu. The final resolution of GS+MgATP/Glu is 1.7Å.

**Figure S7.**
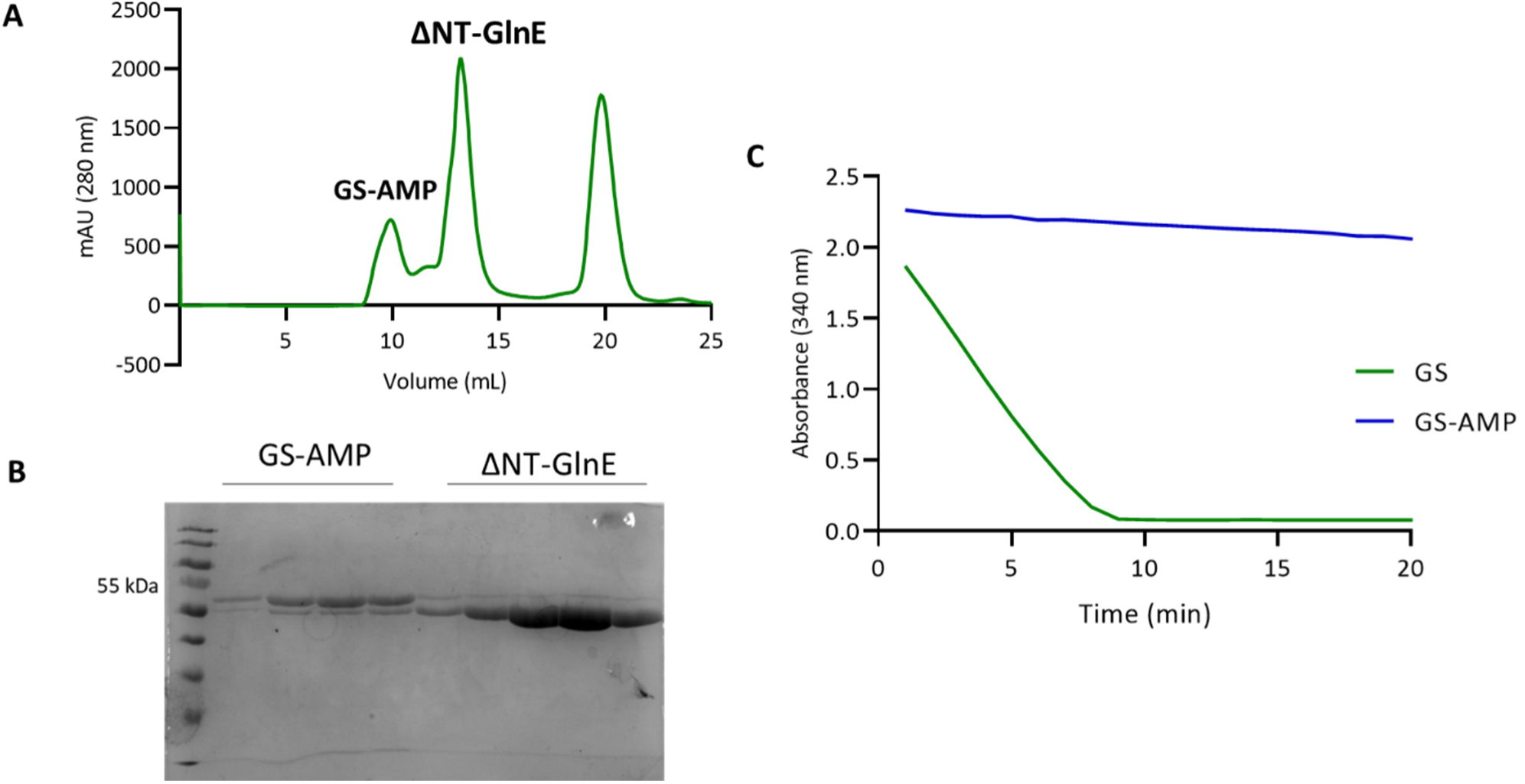
*In vitro* adenylylation of GS for electron cryo-microscopy experiments. A) The SEC chromatogram shows a peak at 9.935 mL, corresponding to the elution of GS as a dodecamer (641 kDa), and a second peak at 13.185 mL, corresponding to the elution of ΔNT-GlnE as a dimer (117 kDa). The third peak corresponds to 3 kDa particles, which are likely an artifact. B) The SDS-PAGE corresponding to the chromatogram peaks indicates that the proteins were mostly separated, although a minor fraction of ΔNT-GlnE co-eluted with GS. C) The GS activity coupled to NADH consumption indicates that the activity of the adenylylated GS was abolished after the *in vitro* reaction, suggesting a high degree of adenylylation in the preparation immediately prior to sample freezing.

**Figure S8.**
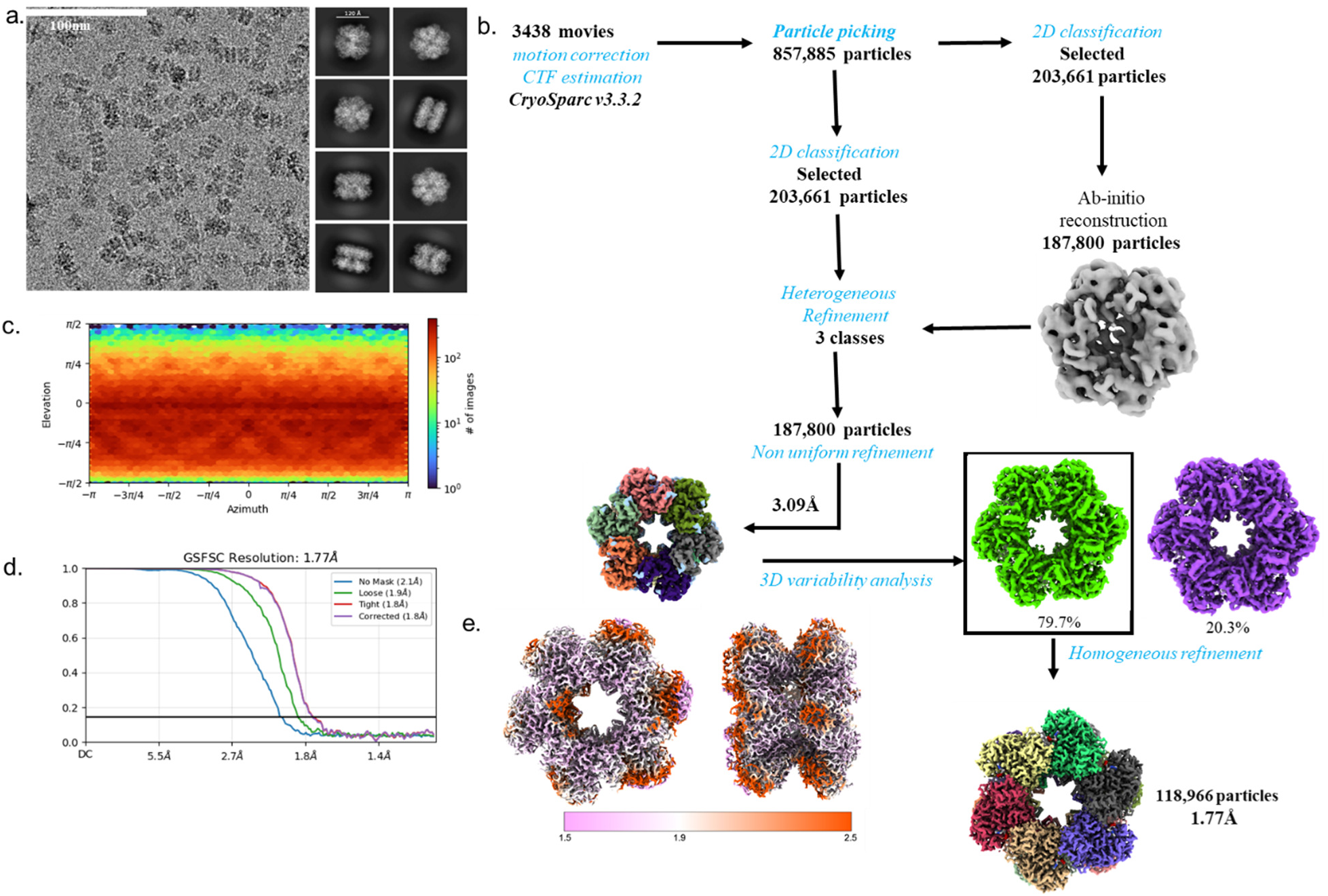
Processing workflow for adenilylated GS in complex with MgATP. A) Representative motion-corrected micrograph showing different orientations of the GS-AMP+MgATP particles and 2D class averages; B) Cryo-EM processing workflow used for obtaining the high-resolution structure of GS-AMP+MgATP. C) Histogram showing the 2D distribution of the particle viewing directions used in the 3D reconstruction: D) Fourier shell correlation (FSC) curves. E) Local resolution maps for different orientations of GS-AMP+MgATP. The final resolution of GS-AMP+MgATP is 1.77 Å.

**Figure S9.**
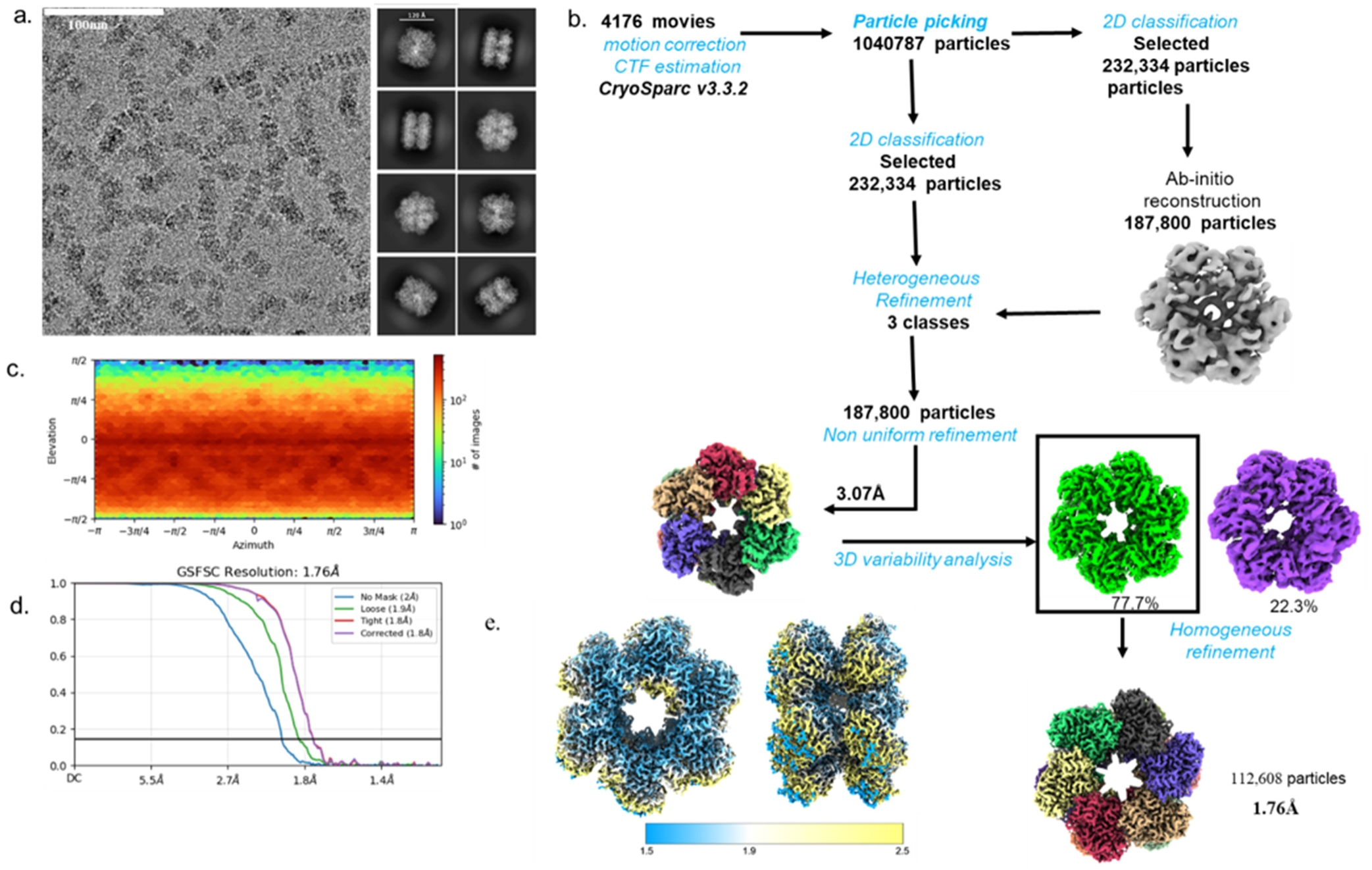
Processing workflow for adenilylated GS in complex with MnADP. A) Representative motion-corrected micrograph showing different orientations of the GS-AMP+MnADP particles and 2D class averages; B) Cryo-EM processing workflow used for obtaining the high-resolution structure of GS-AMP+MnADP. C) Histogram showing the 2D distribution of the particle viewing directions used in the 3D reconstruction: D) Fourier shell correlation (FSC) curves. E) Local resolution maps for different orientations of GS-AMP+MnADP. The final resolution of GS-AMP+MnADP is 1.76 Å.

**Figure S10.**
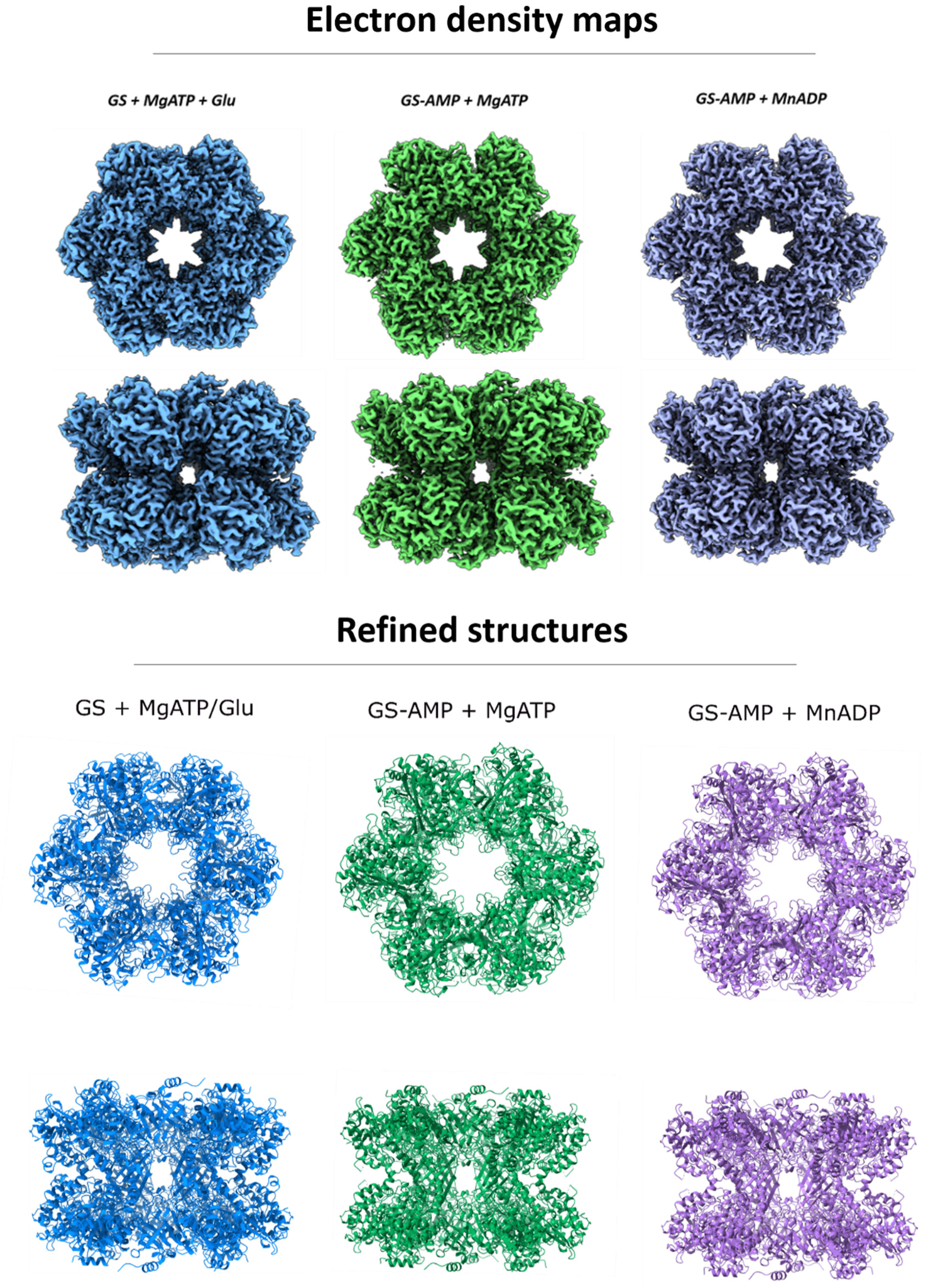
Dodecameric assembly of GS structures solved. All the three structures solved exhibited a dodecameric assembly as found to other GS Type 1. The dodecamer is formed by two hexameric rings joint face-to-face, with a central pore. The AMPylation did not change the overall assembly structure. Blue: unmodified GS in complex with MgATP and Glu; Green: GS-AMP in complex with MgATP; Purple: GS-AMP in complex with MnADP.

**Figure S11.**
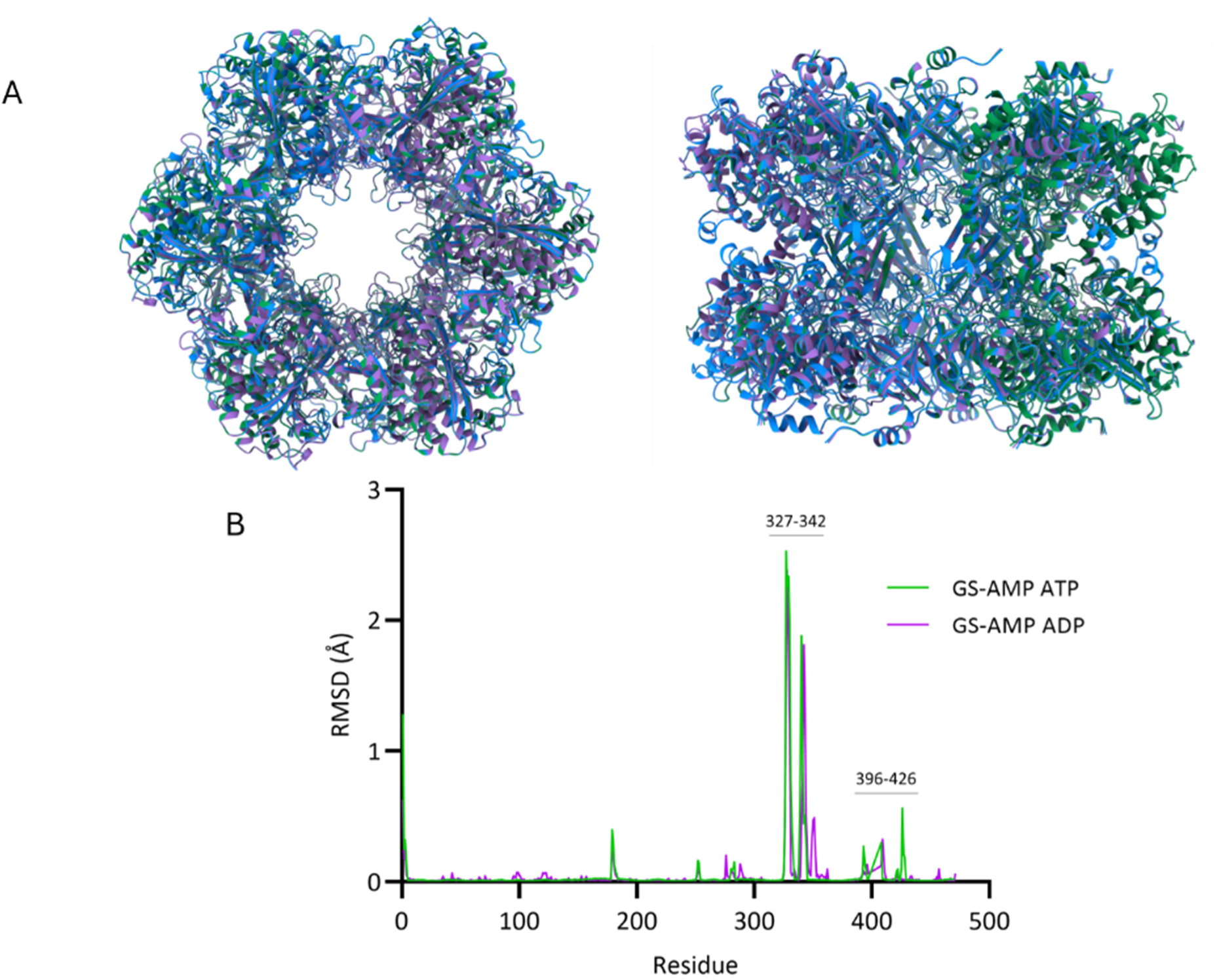
Superposition of all solved GS structures. A) Structural alignment of unmodified and adenylylated GS in complex with MgATP and MnADP. The RMSD values were determined to be 0.378 Å and 0.282 Å between unmodified GS and GS-AMP+MgATP and GS-AMP+MnATP, respectively, suggesting that AMPylation does not significantly change the overall arrangement of the GS structure. B) The major structural deviations were observed in segments 327–342 and 396–426, which harbor the catalytic Arg342 and the AMP-loop, respectively. Blue: GS+MgATP/Glu; Green: GS-AMP+MgATP; Purple: GS-AMP+MnADP.

**Figure S12.**
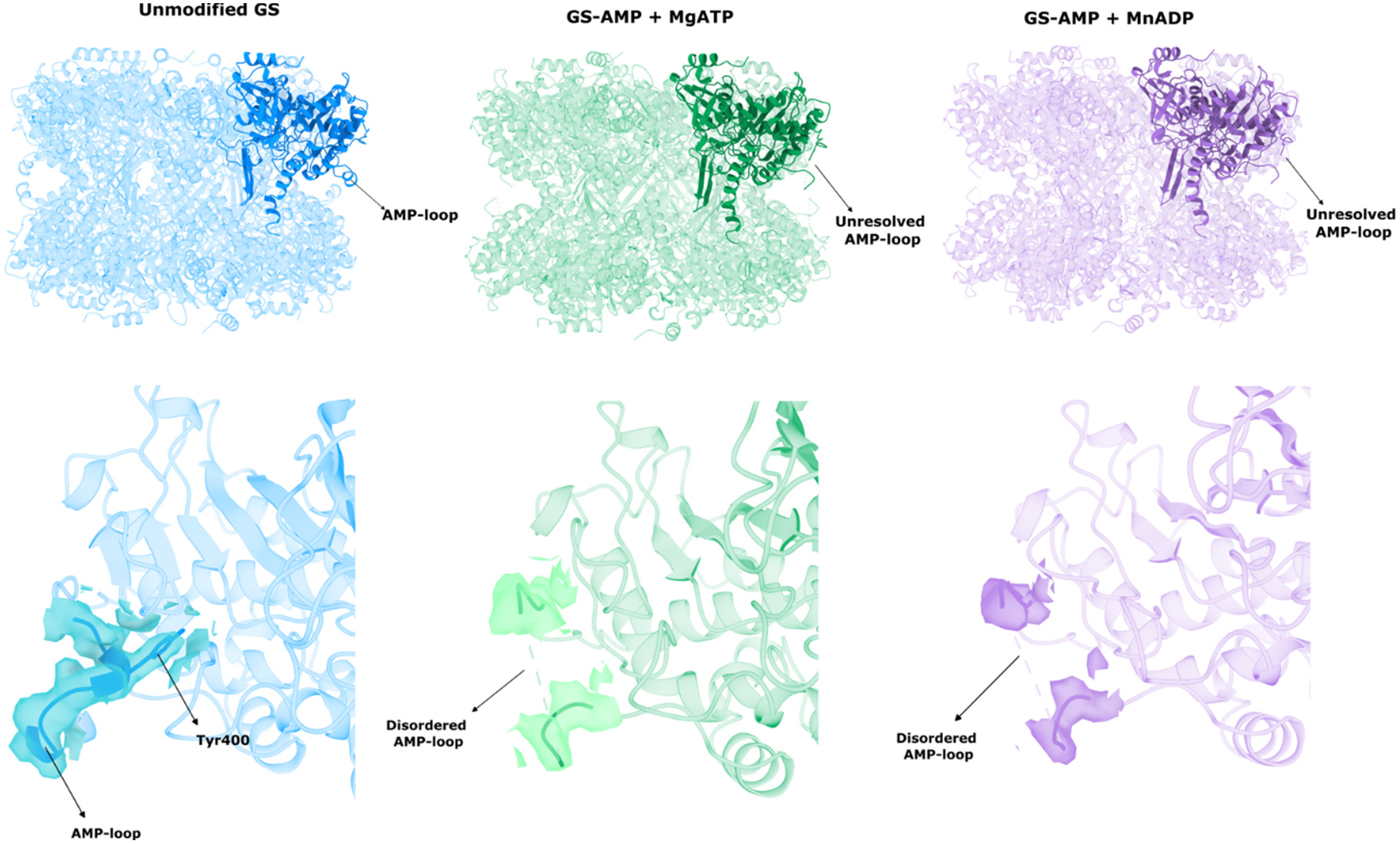
AMP-loop electron density. The AMP-loop electron density was determined only for the unmodified GS, but was not solved for the adenylylated ones, suggesting that the attachment of the AMP moiety renders the AMP-loop flexible. Blue: GS+MgATP/Glu; Green: GS-AMP+MgATP; Purple: GS-AMP+MnADP.

**Figure S13.**
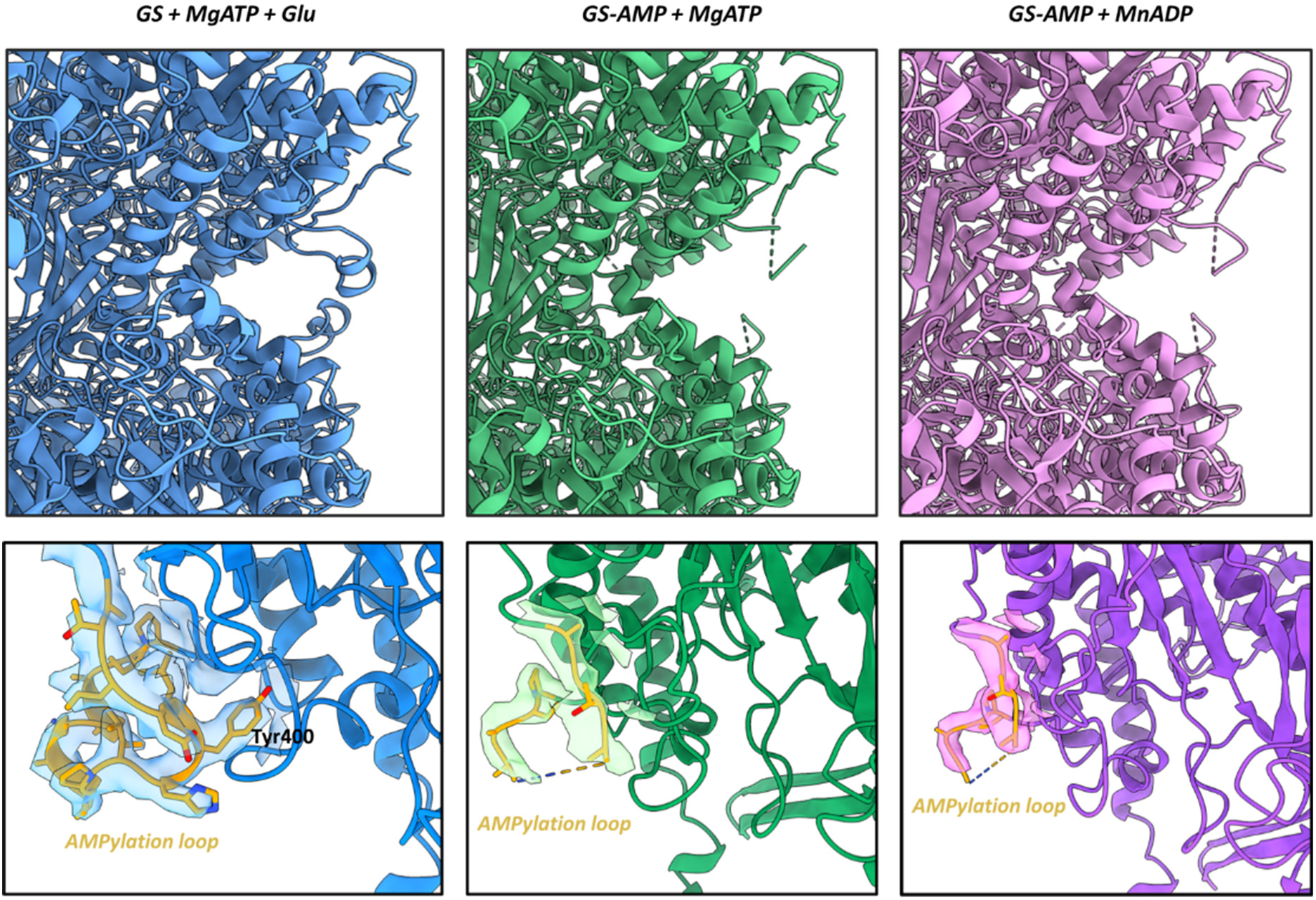
AMP effect on the AMP-loop. Electron density for the AMP-loop was exclusively observed in the unmodified GS structure, remaining unresolvable in the adenylylated forms. This indicates that attachment of the AMP moiety increases the conformational flexibility of the AMP-loop. Blue: GS+MgATP/Glu; Green: GS-AMP+MgATP; Purple: GS-AMP+MnADP.

**Figure S14.**
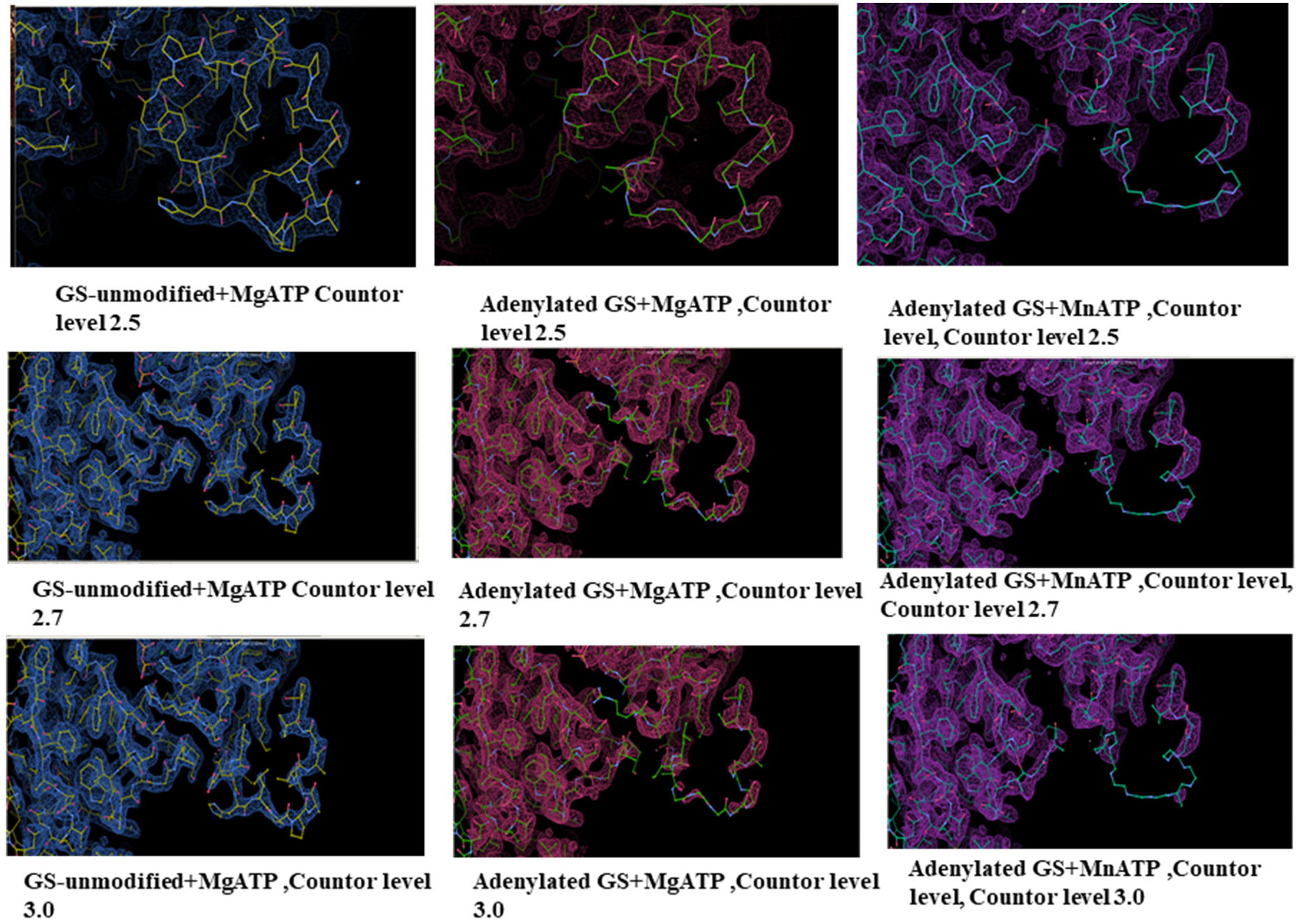
AMP effect on the AMP-loop at diferente countour levels. Electron density for the AMP-loop was observed in the unmodified GS structure, remaining unresolvable in the adenylylated forms, since in higher countour lavels the electon densities were lost. This indicates that attachment of the AMP moiety increases the conformational flexibility of the AMP-loop. Blue: GS+MgATP/Glu; Green: GS-AMP+MgATP; Purple: GS-AMP+MnADP.

**Figure S15.**
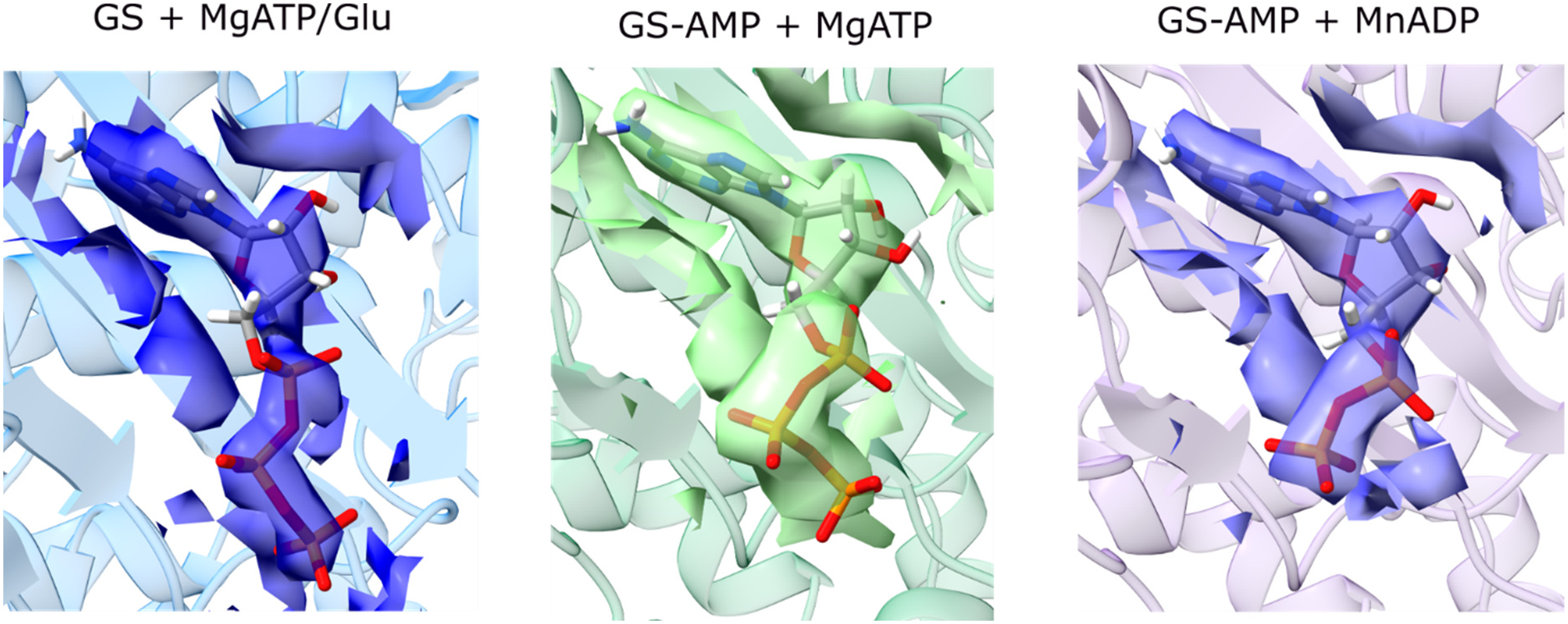
Nucleotide electron densities. The electron densities for the nucleotides ATP and ADP were determined for all three structures, allowing the modeling of the molecules in the nucleotide-binding site. In the unmodified GS, the ATP γ-phosphate is positioned toward the center of the active site, while in the adenylylated structures, the phosphates are far from the center of the active site. This differential position is supposed to be responsible for the inhibition of catalytic activity.

**Figure S16.**
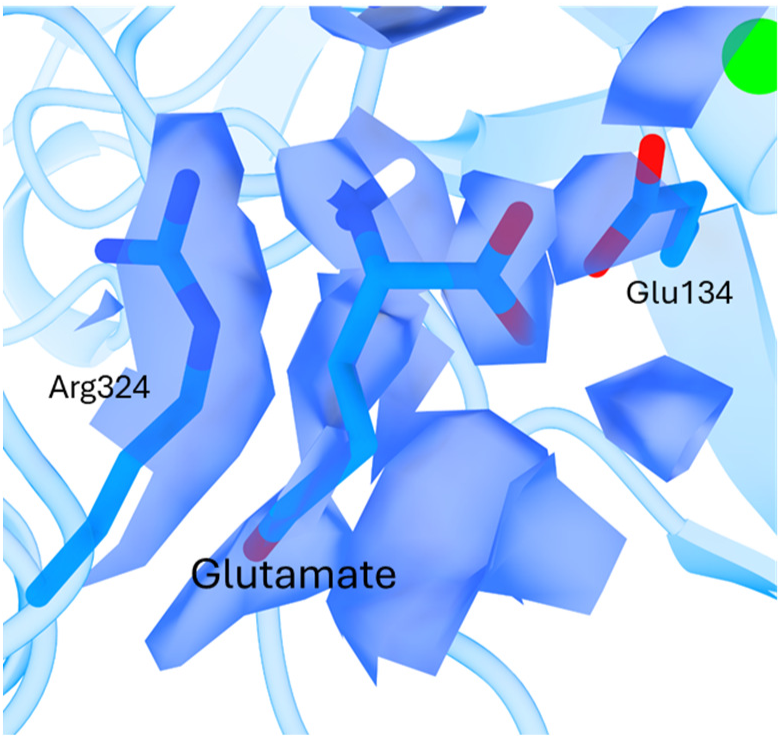
Non-productive glutamate orientation. Electron density for glutamate was resolved in unmodified GS in a non-productive conformation, with its side chain directed away from the active site center. This suggests that the binding site can accommodate glutamate in an orientation distinct from the catalytic conformation, indicating that substrate repositioning must occur for the reaction to take place.

**Table S1.** Cryo-EM data collection, refinement and validation statistics.

|  | *H.seropedicae*<br>GS+MgATP+Glu | *H. seropedicae*<br>GS-AMP+MnADP | *H.seropedicae*<br>GS-AMP+MgATP |
| --- | --- | --- | --- |
| **Data collection and processing** |  |  |  |
| Magnification | 215k | 215 k | 215 k |
| Voltage (kV) | 300 | 300 | 300 |
| Electron exposure (e<sup>-</sup>/Å<sup>2</sup>) | 40 | 40 | 40 |
| Defocus range (μm) | -0.5 to -2.0 | -0.5 to -2.0 | -0.5 to -2.0 |
| Pixel size (Å) | 0.572 | 0.572 | 0.572 |
| Symmetry imposed | *D*6 | *D*6 | *D*6 |
| Initial particle images (no.) | 2,795,951 | 1,040,787 | 857,885 |
| Final particle images (no.) | 376,777 | 112,608 | 118,966 |
| Map resolution (Å) | 1.71 | 1.76 | 1.77 |
| FSC threshold | 0.143 | 0.143 | 0.143 |
| Map resolution range (Å) | 1.5 -2.3 | 1.5-2.5 | 1.5-2.5 |
| **Refinement** |  |  |  |
| Initial model used (PDB code) | de novo, AlphaFold | GS+MgATP | GS+MgATP |
| Map/Model resolution (Å) | 1.78 | 1.82 | 1.87 |
| FSC threshold | 0.5 | 0.5 | 0.5 |
| Map sharpening B factor (Å<sup>2</sup>) | n/a | n/a | n/a |
| Model composition | 44,122 | 42,126 | 42,608 |
| Non-hydrogen atoms | 5,664 | 5531 | 5,530 |
| Protein residues | 12 | 0 | 12 |
| Ligands |  |  |  |
| Mg<sup>2+</sup> | 24 | 0 | 24 |
| Mn<sup>2+</sup> | 0 | 24 | 0 |
| ADP | 0 | 12 | 0 |
| Glu | 12 | 0 | 0 |
| ATP | 12 | 0 | 12 |
| B factors (Å<sup>2</sup>) | 104.89 | 92.07 | 106.59 |
| Protein | 89.55 | 80.92 | 81.39 |
| Ligand |  |  |  |
| R.m.s. deviations | 0.002 | 0.002 | 0.002 |
| Bond lengths (Å) | 0.475 | 0.509 | 0.468 |
| Bond angles (°) |  |  |  |
| Validation | 1.51 | 1.59 | 1.45 |
| MolProbity score Clashscore | 7.44 | 9.14 | 8.38 |
| Poor rotamers (%) | 0.57 | 0.46 | 0.33 |
| Ramachandran plot | 97.48 | 97.50 | 98.07 |
| Favored (%) | 2.52 | 2.43 | 1.92 |
| Allowed (%) | 0.0 | 0.07 | 0.02 |
| Disallowed (%) |  |  |  |

